# Apically localized PANX1 impacts neuroepithelial expansion in human cerebral organoids

**DOI:** 10.1101/2023.07.28.550996

**Authors:** Rebecca J. Noort, Robert T. Flemmer, Craig S. Moore, Thomas J. Belbin, Jessica L. Esseltine

## Abstract

Dysfunctional paracrine signaling through Pannexin 1 (PANX1) channels is linked to several adult neurological pathologies and emerging evidence suggests that PANX1 plays an important role in human brain development. It remains unclear how early PANX1 influences brain development, or how loss of PANX1 alters the developing human brain. Using a cerebral organoid model of early human brain development, we find that PANX1 is expressed at all stages of organoid development from neural induction through to neuroepithelial expansion and maturation. Interestingly, PANX1 cellular distribution and subcellular localization changes dramatically throughout cerebral organoid development. During neural induction, PANX1 becomes concentrated at the apical membrane domain of neural rosettes where it co-localizes with several apical membrane adhesion molecules. During neuroepithelial expansion, *PANX1*-/- organoids are significantly smaller than control and exhibit significant gene expression changes related to cell adhesion, Wnt signaling and non-coding RNAs. As cerebral organoids mature, PANX1 expression is significantly upregulated and is primarily localized to neuronal populations outside of the ventricular-like zones. Ultimately, PANX1 protein can be detected in all layers of a 21-22 post conception week human fetal cerebral cortex. Together, these results show that PANX1 is dynamically expressed by numerous cell types throughout embryonic and early fetal stages of human corticogenesis and loss of PANX1 compromises neuroepithelial expansion due to dysregulation of cell-cell and cell-matrix adhesion, perturbed intracellular signaling, and changes to gene regulation.

## Introduction

Human brain development follows a series of intricately choreographed events involving large cellular migrations and rearrangements, changes in cell morphology, and cell fate specification. These activities are locally organized through exquisite spatial and temporal control of signaling events between neighboring cells. Dysfunctional paracrine signaling through Pannexin 1 (PANX1) channels is linked to several adult neurological pathologies and human germline *PANX1* variants have been associated with severe neurological deficits and autism spectrum disorder (Davis et al., 2012; Shao et al., 2016). Studies in postnatal rodent models reveal PANX1 expression across various neural cell types including neurons, glia, and neural progenitor cells (Zappalà et al., 2007; Bond and Naus, 2014; Seo et al., 2021). In postnatal murine neural precursor cells (NPCs), PANX1 restricts neuronal differentiation by impeding neurite extension and cell migration via the channels’ ATP release functions and interactions with the cytoskeleton (Wicki-Stordeur et al., 2012; Wicki-Stordeur and Swayne, 2013). Others have demonstrated PANX1 localization at neuronal synapses where the channels help to replenish extracellular ATP, negatively regulate dendritic spine density, and maintain synaptic strength (Prochnow et al., 2012; Sanchez-Arias et al., 2020). However, it remains unclear how early PANX1 influences human brain development, or which cell types express PANX1 in the developing human brain.

Recent reports have revealed that PANX1 is expressed in some of the earliest cell types in human development including human oocytes, pluripotent stem cells, and the three embryonic germ layers (definitive endoderm, mesoderm, and ectoderm) (Hainz et al., 2018; Sang et al., 2019; Noort et al., 2021). PANX1 channels are also expressed throughout embryonic brain development. *PANX1* transcript expression is robust in the developing mouse cerebral cortex, cerebellum, and olfactory bulbs where maximum *PANX1* expression occurs at murine embryonic day 18 and declines thereafter (Ray et al., 2005). Gene expression analyses curated by BrainSpan indicate that a similar pattern occurs in the human system as *PANX1* transcript expression in various brain structures is high at 8 post conception weeks (pcw) (earliest timepoint assessed) but diminishes around 26 pcw (Brainspan.org). The Human Protein Atlas reports moderate-to-high PANX1 protein abundance in the adult human cerebral cortex (humanproteinatlas.org). Given this dynamic pattern of PANX1 expression, we expect that PANX1-mediated cellular communication influences proper development of neural tissues.

To date, the cellular and subcellular localization of PANX1 protein throughout human embryonic and early fetal brain development have not been investigated. Here we use iPSC-derived neural precursor cells, neurons, and cerebral organoids to investigate PANX1 expression and localization as iPSCs differentiate to neural cell types and organized cortical structures. Cerebral organoids recapitulate a variety of human brain regions including the cerebral cortex, hippocampus, choroid plexus, and retinal tissue and contain a variety of cell types including neural progenitors (like neuroepithelial cells and radial glia), neurons, astrocytes, oligodendrocytes, retinal pigment epithelial cells, and ependymal cells (Lancaster et al., 2013; Lancaster and Knoblich, 2014; Di Lullo and Kriegstein, 2017; Renner et al., 2017). Importantly, the cells within cerebral organoids self-organize to form cortical-like layers like those seen in the developing human brain (Di Lullo and Kriegstein, 2017) making cerebral organoids a powerful tool to study the embryonic and early fetal stages of human brain development.

Given that PANX1 is expressed in the earliest cell types of human development and is linked to neurological disease, we sought to explore PANX1 expression and localization throughout early stages of human brain development. Immunostaining of a 21-22 pcw (midgestation) human fetal cerebral cortex reveals PANX1 protein expression in all cortical layers, with heightened signal intensity in the marginal zone. We observe concentrated PANX1 expression at the apical membrane domain of neuroepithelial-stage iPSC-derived cerebral organoids whereas more mature organoids exhibit the heaviest PANX1 expression within the emerging neuronal layers. CRISPR-Cas9 *PANX1* gene ablation results in stunted neuroepithelial expansion and dysregulation of genes related to cell signaling, cell adhesion, and expression of non-coding RNAs.

## Materials & Methods

### Induced Pluripotent Stem Cells

These studies were approved by the Newfoundland and Labrador Health Research Ethics board (HREB # 2018.210). A male iPSC line (GM25256) was purchased from the Coriell Institute for Medical Research (Cat# GM25256, Coriell, Camden, NJ, USA). Female iPSCs (43Q) were created as described previously (Esseltine et al., 2017) and obtained through a material transfer agreement with The University of Western Ontario. Both cell lines were derived from fibroblasts of apparently healthy individuals with no known genetic pathologies.

iPSCs were cultured in a humidified 37°C cell culture incubator buffered with atmospheric oxygen and 5% CO_2_. The iPSCs were grown on Geltrex™-coated (Cat# A141330, ThermoFisher, Waltham, MA, USA) culture dishes and fed daily with Essential 8™ medium (Cat# A1517001, ThermoFisher) or mTeSR™ Plus (Cat #05825, STEMCELL Technologies, Vancouver, BC, CAN) maintenance medium. Every 4-5 days, iPSCs were passaged as small aggregates using a cell scraper and 0.5 mM EDTA (Cat #AM9260G, ThermoFisher) prepared in Ca^2+^/Mg^2+^-free phosphate buffered saline (PBS; Cat# 319-005-CL, WISENT Inc., St. Bruno, QC, CAN) (Beers et al., 2012) when the colonies exhibited smooth borders and tight cell packing. Aggregates were seeded into fresh Geltrex™-coated wells containing Essential 8™ or mTeSR™ Plus at split ratios of 1:5 to 1:50. StemPro™ Accutase™ (Cat# A1110501, ThermoFisher) was used to create suspensions of single cell iPSCs. Single cells were plated in medium supplemented with 10 µM of the rho-associated kinase inhibitor (ROCKi), Y-27632 (Cat# 100005583, Cayman Chemicals, Ann Arbor, MI, USA) to promote single cell iPSC survival (Watanabe et al., 2007). After thawing from liquid nitrogen stocks, iPSCs were maintained in culture for up to 20 weeks at which point a new vial was thawed. Evaluation of our iPSC cell banks with the hPSC Genetic Analysis Kit (Cat # 07550, STEMCELL Technologies) confirmed normal copy number at various mutation hotspots and assessment with a Mycoplasma PCR Detection Kit (Cat# G238, Applied Biological Materials Inc., Richmond, BC, CAN) indicated that cell stocks are free of mycoplasma.

### Monolayer Differentiation to Neural Progenitors and Neurons

Human iPSCs were differentiated to neural progenitor cells according to the methodology described by (Yan et al., 2013) with several modifications. On day 0, singularized iPSCs were plated at a density of 200,000 viable cells/cm^2^ onto Geltrex™-coated dishes containing Gibco™ PSC Neural Induction Medium (Cat# A1647801, ThermoFisher) supplemented with 10 µM ROCKi. Daily feeds with PSC Neural Induction Medium without ROCKi were administered until day 7 when the cells were singularized and re-plated at a density of 200,000 viable cells/cm^2^ onto Geltrex™-coated dishes containing Neural Stem Cell (NSC) Expansion Medium supplemented with 10 µM ROCKi. NSC Expansion Medium consists of 49% Neurobasal (Cat# 21103049, ThermoFisher), 49% Advanced DMEM/F12 media (Cat# 12634010, ThermoFisher), and 2% (1X) Neural Induction Supplement (Cat# A16477-01, ThermoFisher). Cells were fed daily with NSC Expansion Medium without ROCKi and seeded into new Geltrex™-coated wells every 7 days. On day 21 or 22 the resultant NPCs were assayed or differentiated further to neurons.

For differentiation to neurons, day 21 or 22 NPCs were passaged as single cells and seeded at 50,000 cells/cm^2^ onto culture wells coated with 10 µg/mL laminin (Cat# 354232, Corning Inc, Corning, NY, USA) containing Neuron Differentiation Medium supplemented with 10 µM ROCKi. Neuronal Differentiation Medium consists of ∼96% Neurobasal (Cat # 21103049, ThermoFisher), 2% (1X) B-27 (Cat #17504044, ThermoFisher), 1% (1X) non-essential amino acids (Cat# 321-011-EL, WISENT), 20 ng/mL brain-derived neurotropic factor (BDNF; Cat# 78005, STEMCELL Technologies), 20 ng/mL glial cell-derived neurotropic factor (GDNF; Cat# 78058, STEMCELL Technologies), and 200 µM L-ascorbic acid 2-phosphate sesquimagenesium salt hydrate (Cat# A8960, MilliporeSigma, Burlington, MA, USA). Half medium changes with Neuron Differentiation Medium were performed every other day for 14 days.

### Cerebral Organoids

Cerebral organoids were generated using the STEMdiff™ Cerebral Organoid Kit and STEMdiff™ Cerebral Organoid Maturation Kit (Cat# 08570 & 08571, STEMCELL Technologies) according to the manufacturer’s instructions with the following modifications: On Day 0, 96-well round-bottom plates (Cat# 351177, Corning) were rinsed with a solution of 5% Pluronic™ F-127 (Cat# P2443, MilliporeSigma) prepared in deionized water to confer an anti-adherent coating (Kurosawa, 2007). On Day 7 the organoids were subjected to high throughput Geltrex™ embedding in Expansion Medium according to Chew et al., with slight modification (Chew et al., 2022). Briefly, ice cold liquid Geltrex™ was added at 1:50 dilution to ice-cold Expansion Medium. The organoids were quickly transferred into the cold Expansion Medium with Geltrex™ and re-plated into a fresh Pluronic™ F-127-coated 6-well dish (Cat# 140685, ThermoFisher).

### Human Fetal Brain Preparation

These studies were approved by the Newfoundland and Labrador Health Research Ethics board (HREB # 2014.216. Formalin fixed and paraffin embedded samples from a 21-22 pcw human fetal brain were cut to a thickness of 5 µm using a microtome and deposited onto positively charged glass slides (Cat# ER4951PLUS, FisherScientific). The sections were dewaxed with xylene substitute (MilliporeSigma, Cat# 78475) and rehydrated with graded ethanol solutions. After rehydration, the sections were subjected to antigen retrieval and antibody staining for immunofluorescence.

### Immunofluorescence imaging

Monolayer cultures grown on Geltrex™ or laminin-coated #1.5 glass coverslips were fixed in 10% buffered formalin (Cat# CA71007-344, VWR, Radnor, PA, USA) for 10 minutes at room temperature and permeabilized with PBS-T (Ca^2+^/Mg^2+^-free PBS + 0.1% TWEEN® 20 (Cat# BP337-500, FisherScientific, Waltham, MA, USA)) for 20 minutes followed by 0.1% Triton™ X-100 (Cat# T5832, MilliporeSigma) in Ca^2+^/Mg^2+^-free PBS + for 10 minutes. Samples were incubated overnight at 4°C in primary antibodies diluted in PBS-T with 3% bovine serum albumin (BSA; Cat# 800-095-EL, WISENT Inc.) and 0.1% NaN_3_ according to Table 1. Secondary antibodies and/or dyes (Table 1) prepared in PBS-T were applied for 2 hours at room temperature. All Alexa Fluor® and HRP (horseradish peroxidase) conjugated secondary antibodies were purchased from ThermoFisher. Slides were mounted using Mowial®488 reagent with 1,4-diazabicyclo[2.2.2]octane (DABCO) antifade compound according to the formulation described by Cold Spring Harbour (Laboratory, 2007). For whole-mount imaging, fixed organoids were permeabilized and stained according to the methodology described above and transferred to an 8-well µ-slide high-end microscopy chamber slide (Cat# 80826, ibidi, Gräfelfing, DEU) for confocal imaging.

**Table 1.**
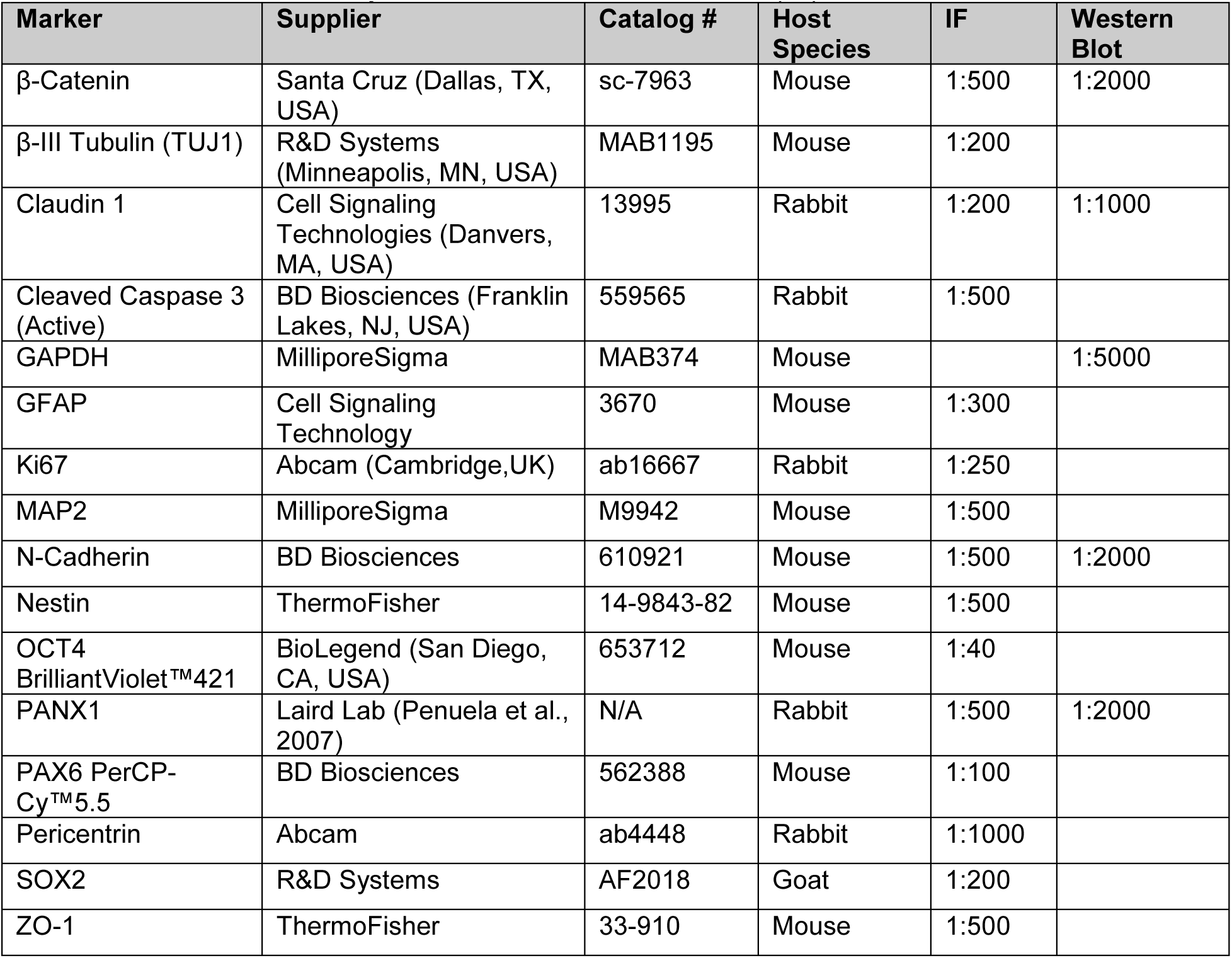
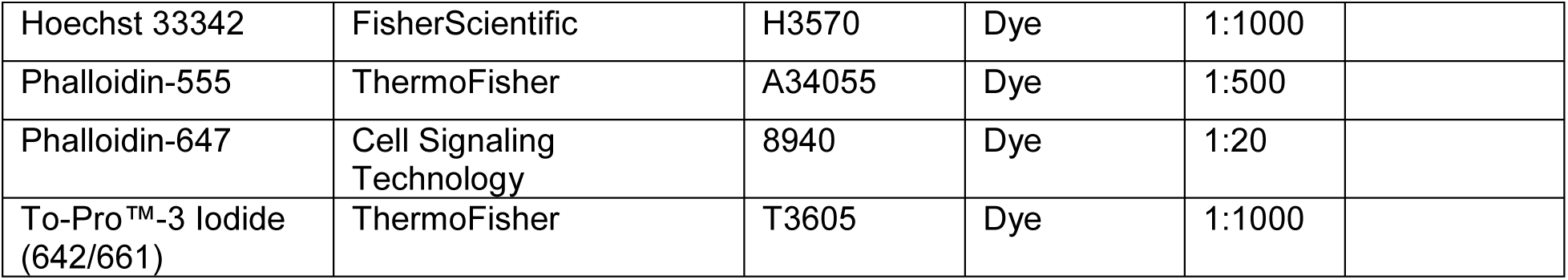
Antibodies and Dyes for Immunofluorescence (IF) and Western Blot.

| Marker | Supplier | Catalog # | Host Species | IF | Western Blot |
| --- | --- | --- | --- | --- | --- |
| β-Catenin | Santa Cruz (Dallas, TX, USA) | sc-7963 | Mouse | 1:500 | 1:2000 |
| β-III Tubulin (TUJ1) | R&D Systems (Minneapolis, MN, USA) | MAB1195 | Mouse | 1:200 | |
| Claudin 1 | Cell Signaling Technologies (Danvers, MA, USA) | 13995 | Rabbit | 1:200 | 1:1000 |
| Cleaved Caspase 3 (Active) | BD Biosciences (Franklin Lakes, NJ, USA) | 559565 | Rabbit | 1:500 |  |
| GAPDH | MilliporeSigma | MAB374 | Mouse |  | 1:5000 |
| GFAP | Cell Signaling Technology | 3670 | Mouse | 1:300 |  |
| Ki67 | Abcam (Cambridge,UK) | ab16667 | Rabbit | 1:250 |  |
| MAP2 | MilliporeSigma | M9942 | Mouse | 1:500 |  |
| N-Cadherin | BD Biosciences | 610921 | Mouse | 1:500 | 1:2000 |
| Nestin | ThermoFisher | 14-9843-82 | Mouse | 1:500 |  |
| OCT4 BrilliantViolet™421 | BioLegend (San Diego, CA, USA) | 653712 | Mouse | 1:40 |  |
| PANX1 | Laird Lab (Penuela et al., 2007) | N/A | Rabbit | 1:500 | 1:2000 |
| PAX6 PerCP-Cy™5.5 | BD Biosciences | 562388 | Mouse | 1:100 |  |
| Pericentrin | Abcam | ab4448 | Rabbit | 1:1000 |  |
| SOX2 | R&D Systems | AF2018 | Goat | 1:200 |  |
| ZO-1 | ThermoFisher | 33-910 | Mouse | 1:500 |  |
| Hoechst 33342 | FisherScientific | H3570 | Dye | 1:1000 |  |
| Phalloidin-555 | ThermoFisher | A34055 | Dye | 1:500 |  |
| Phalloidin-647 | Cell Signaling Technology | 8940 | Dye | 1:20 |  |
| To-Pro™-3 Iodide (642/661) | ThermoFisher | T3605 | Dye | 1:1000 |  |

Cerebral organoids were fixed overnight (∼20 hours) in 10% normal buffered formalin and cryogenically prepared according to the methodology described in STEMCELL Technologies’ Document #27171, Version 1.0.0, Nov 2019. Briefly, organoids were first dehydrated in Ca^2+^/Mg^2+^-free PBS supplemented with 30% sucrose for 1-4 days at 4°C until the organoids sank. Dehydrated organoids were then incubated for 1 hour at 37°C in gelatin embedding solution consisting of 10% sucrose and 7.5% gelatin (Cat# G1890, MilliporeSigma) prepared in Ca^2+^/Mg^2+^-free PBS. The organoids were then snap frozen in a slurry of dry ice and isopentane followed by cryosectioning at thickness of 14 µm and deposition onto positively charged glass microscope slides (Cat# ER4951PLUS, FisherScientific). For antigen retrieval, sections were placed into a plastic container with pH 6.0 citrate buffer: 0.294% Tri-sodium citrate (dihydrate) (Cat# A12274, Alfa Aesar, Tewksbury, MA, USA) + 0.05% TWEEN® 20 and heated in a food steamer (Hamilton Beach, Glen Allen, VA, USA) for 20 minutes. Immunostaining and mounting were performed as stated above with antibodies and dyes listed in Table 1.

### Phase contrast imaging

Phase contrast images of monolayer cells and organoids were taken on a Zeiss AxioObserver microscope using 5X/0.12 NA A-Plan and 10X/0.25 NA Ph1 objectives. Images from these microscopes were taken in 8-bit greyscale using an Axiocam MRm camera and AxioVision Version 4.8.2 software. All phase contrast imaging equipment is from Carl Zeiss Microscopy (Jena, DEU).

### Organoid size measurements

Area measurements from phase contrast images of day 10 cerebral organoids were performed automatically using the batch macro code in FIJI open source software (Schindelin et al., 2012). Area measurements from images that contained debris (fibers and unincorporated cells) were performed manually by tracing around the object’s periphery and excluding debris protuberances. The macro shown here computes object area for entire folders of phase contrast images that were taken on the same microscope, at the same magnification. The macro can be adjusted for different magnifications and microscopes by changing the parameters in “Set Scale”.

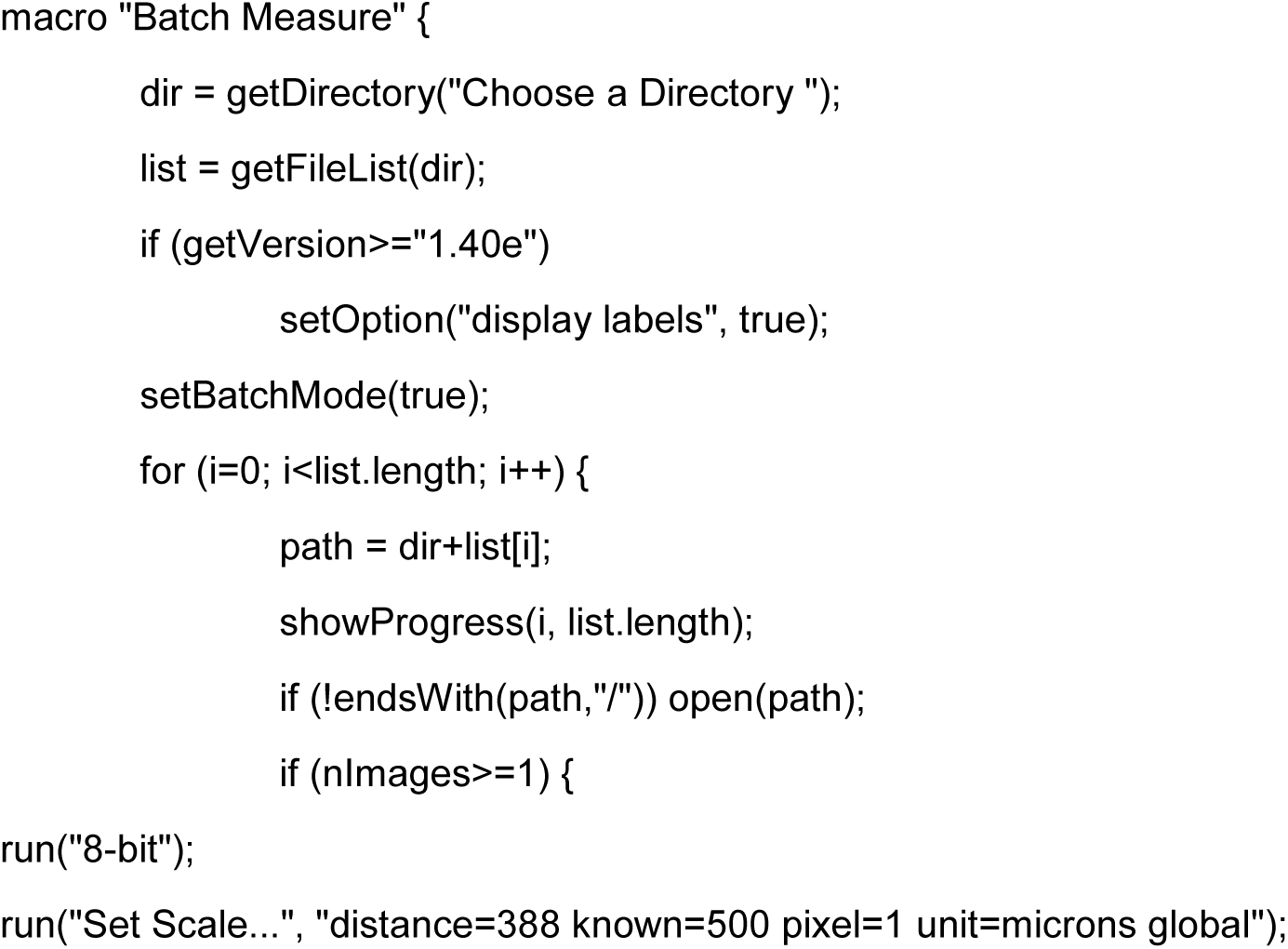
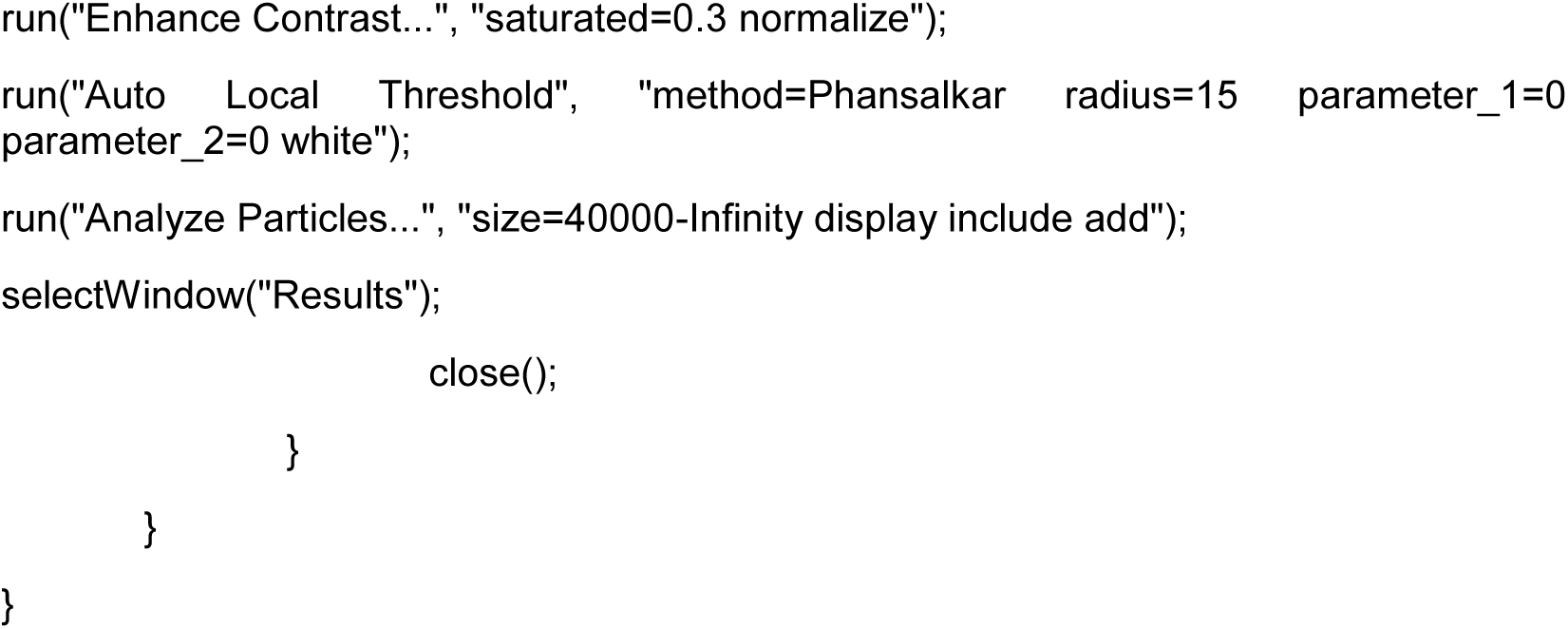

### Confocal Microscopy and Image Analysis

Fluorescent confocal images were primarily acquired on an Olympus Fluoview FV10i—W3 confocal microscope (Olympus, Tokyo, JPN) fitted with a 10X/0.4 NA or 60X/1.2 NA lens and Fluoview version 2.1.17 software. The following lasers were used to visualize fluorophores: Hoechst/Brilliant Violet™ 421 (405 nm laser); Alexa Fluor® 488 (473 nm laser); Alexa Fluor® 555/Phalloidin-555 (559 nm laser); Alexa Fluor® 647/Phalloidin-647/To-pro™-3 iodide (635 nm laser). Additional images were taken on an Olympus FV1000 confocal microscope fitted with 10X/0.4 NA, 20X/0.75 NA, 40X/0.95 NA or 60X/1.42 NA objectives and the following lasers: 405 nm, 458 nm, 568 nm, and 633 nm. Tiled images of 21-22 pcw human cerebral cortex were taken on a ZEISS LSM 900 with Airyscan 2 fitted with 20X/0.8NA and the following lasers: 405 nm, 488 nm, 561 nm, 640 nm. Images were analyzed using FIJI where fluorescent confocal images were occasionally subjected to equivalent brightness/contrast enhancement to improve image clarity.

We used Manders’ colocalization coefficients to describe the fraction of PANX1 colocalizing with a second target (Dunn et al., 2011; Boassa et al., 2014). Manders’ colocalization coefficient values range from 0.0 to 1.0 where values of 0.0 signify no pixel overlap and values of 1.0 denote identical spatial occupation between two signals (Dunn et al., 2011). We performed colocalization analysis in FIJI using the JACoP plugin with Coste’s automatic thresholding (Bolte and Cordelières, 2006). We report the standard error of the mean for Manders’ colocalization coefficients as indicated in the figure legends.

### Whole transcriptome analysis of PANX1 knockout using RNA sequencing

RNA was extracted using the PureLink™ RNA isolation kit (Cat # 12183018A, ThermoFisher) with on column DNase I digestion (Cat# 12185010, ThermoFisher) according to the manufacturers’ instructions. Purified RNA was quantified using a NanoDrop™ 2000 spectrophotometer (Cat# ND-2000, ThermoFisher), and stored at - 80°C until use. High quality RNA was identified by a ʎ260/280 of ≥ 2.0 and ʎ260/230 of ≥ 2.0. RNA was converted into complementary DNA (cDNA) using the High-Capacity cDNA Reverse Transcription Kit (Cat# 4368814, ThermoFisher) according to the manufacturer’s instructions. Typically, 500 ng of RNA were used per 20 µL cDNA reaction. The resulting cDNA was stored at −30°C until use.

Whole transcriptome analysis of gene expression differences in PANX1 knockout cells was carried out by RNA sequencing on a Illumina NovaSeq 6000 S4 PE100 (Genome Quebec). Paired end 100bp reads were assessed for quality control using FastQC (version 0.11.9) (Wingett and Andrews, 2018). Reads were aligned to the Human hg38 reference genome using RNA-Star (Galaxy Version 2.7.8a) with default settings (Dobin et al., 2013) and transcripts were counted using featureCounts (Galaxy version 2.0.1) (Heo et al., 2013). Differential expression of genes between control and *PANX1-/-* cells were based on a model using the negative binomial distribution with DeSeq2 (Galaxy Version 2.11.40.7), with a Benjamini-Hochberg adjusted p-value of less than 0.05 (Love et al., 2014).

Identification of overrepresented groups of genes was carried out using GOseq (Galaxy Version 1.44.0) (Young et al., 2010). The three Gene Ontology (GO) categories were GO:MF (Molecular Function), GO:CC (Cellular Component), GO:BP (Biological Process). Distributions of the numbers of members of a category amongst the differentially expressed genes were determined by the Wallenius non-central hypergeometric distribution. P-values for over representation of the GO term in the differentially expressed genes were adjusted for multiple testing with the Benjamini-Hochberg procedure. GOseq was similarly applied for KEGG (Kyoto Encyclopedia of Genes and Genomes) pathway-based enrichment of differentially expressed genes.

### SDS-PAGE & Western Blot

Cells were lysed with a solution comprising 50 mM Tris-HCl pH 8, 150 mM NaCl, 0.02% NaN_3_, 0.1% Triton™ X-100, 1 mM NaVO_4_, 10 mM NaF, 2 µg/mL leupeptin, and 2 µg/mL aprotinin. Soluble proteins were separated using SDS-PAGE and transferred to a 0.45 µm nitrocellulose membrane (Cat# 1620115, Bio-Rad, Hercules, CA, USA). Primary antibodies (Table 1) were prepared in TBST (15.23 mM Tris HCl, 4.62 mM Tris Base, 150 mM NaCl, and 0.1% TWEEN® 20, adjusted to pH 7.6) + 3% BSA and incubated overnight at 4°C. Secondary antibodies conjugated to HRP were prepared in TBST + 3% BSA and incubated for 1 hour at room temperature. Proteins were visualized with Bio-Rad Clarity™ Western ECL Substrate (Cat# 1705061, Bio-Rad) using a ChemiDoc™ Imaging System (Cat# 12003153, Bio-Rad).

### Statistics

Statistical analyses were performed in GraphPad PRISM Version 6.07. Error bars depict ± standard error of the mean (SEM) when n ≥ 3 biological replicates (independent experiments) unless otherwise stated. Statistical significance for comparisons between 2 groups was determined by unpaired Student’s *t*-test. Statistical significance for comparisons between 3 or more groups was determined by Analysis of Variance (ANOVA) followed by a Tukey’s multiple comparisons test unless otherwise indicated. A p value less than 0.05 is considered statistically significant. * p < 0.05, ** p <0.01, *** p < 0.001.

## Results

### PANX1 is expressed across the human fetal cerebral cortex

The PANX1 literature heavily favors perinatal or postnatal mouse systems. However, the Allen Institute’s Brainspan prenatal laser microdissection (LMD) microarray dataset depicts *PANX1* transcript expression in 21 pcw human fetal brains including cortical regions such as the ventricular zone (VZ), subventricular zone (SVZ), intermediate zone (IZ), subplate (SP), cortical plate (CP), and marginal zone (MZ) (Brainspan.org). To confirm whether PANX1 protein is also expressed in these developing human tissue layers, we performed immunofluorescence confocal imaging on cortical samples from a 21-22 pcw human fetal brain (Figure 1). We find PANX1 signal across all layers of the developing human cerebral cortex with widespread staining throughout the SVZ and brighter manifestation in the marginal zone (Figure 1A). PANX1 signal intensity is diminished in regions with tightly packed nuclei, such as the cortical plate (Figure 1A). In contrast, cortical layers with fewer nuclei such as the marginal zone and subplate display widespread PANX1 staining, concentrated throughout the many processes of MAP2-positive neurons (Figure 1A, B). Interestingly, while we did see some evidence of PANX1 expression within the SOX2-positive stem cells lining the ventricular zone, this staining was much diminished compared to the more mature neuronal layers within the human fetal cortex (Figure 1C). Collectively, we find that PANX1 protein expression is apparent in all cortical layers of the early fetal human brain.

**Figure 1.**
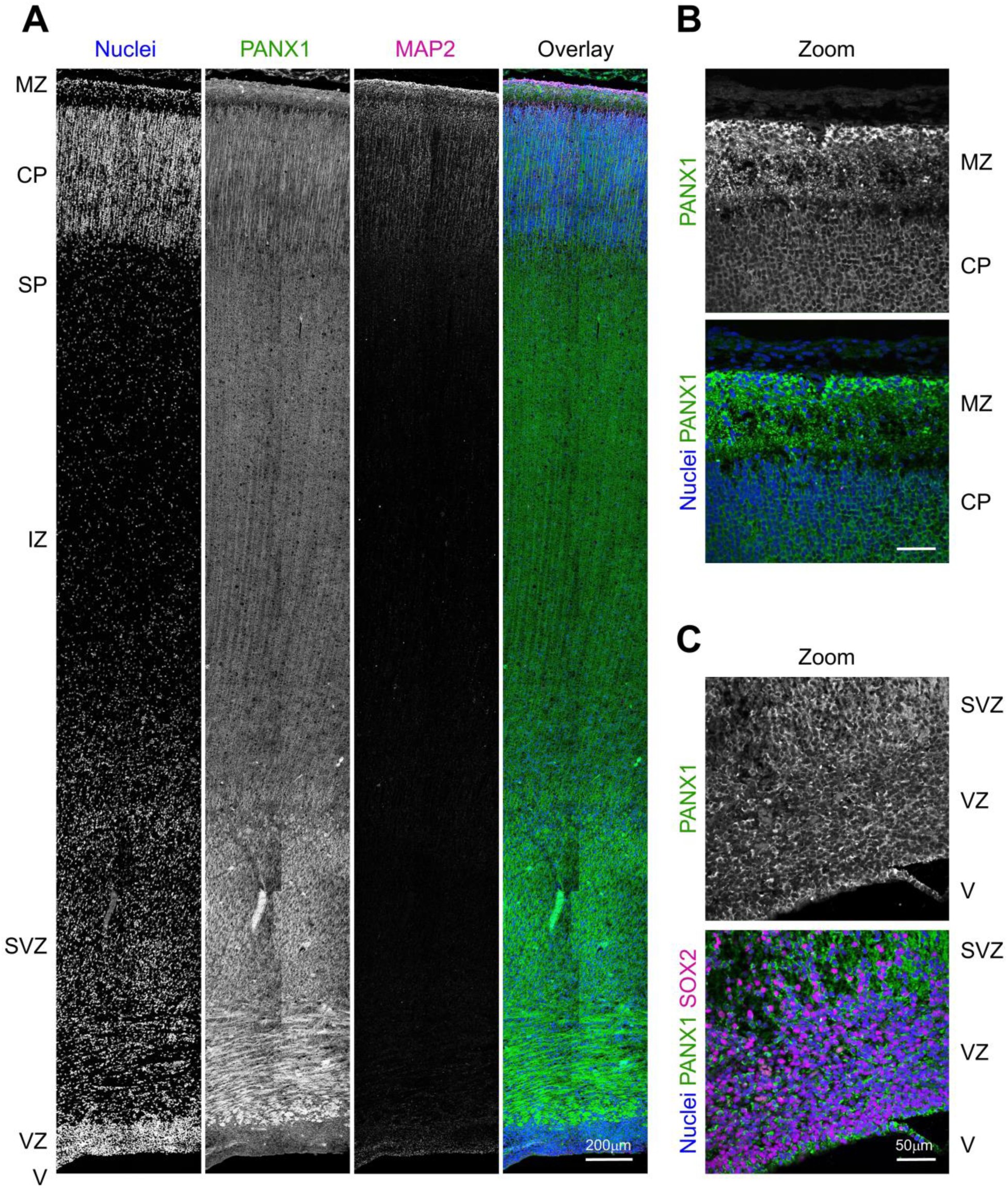
PANX1 protein is expressed across all layers of the human fetal cerebral cortex. **(A)** Representative immunofluorescence confocal micrograph of PANX1 (green) along with MAP2 (magenta) across the span of a 21-22 week (midgestation) human fetal cerebral cortex. **(B, C)** Higher resolution confocal micrographs demonstrating PANX1 cellular distribution in **(B)** the outer cortical layers and **(C)** the subventricular zone. Nuclei (Hoechst, blue). Scale bars as indicated. MZ = marginal zone; CP = cortical plate; SP = subplate; IZ = intermediate zone; SVZ = subventricular zone; VZ = ventricular zone; V = ventricle

### PANX1 is upregulated in neural progenitor cells and neurons compared to undifferentiated iPSCs

The first step to determining how PANX1 influences human brain development is to uncover when and where PANX1 is expressed in the developing human brain. However, human fetal brain samples are precious and few, and a lot of development has already occurred even at the 21-22 pcw timepoint presented in Figure 1. Therefore, once we confirmed PANX1 expression in a 21-22 pcw human fetal cortex, we evaluated PANX1 expression and localization in human iPSCs *in vitro*, and after differentiation into neural precursor cells (NPCs) and mature neurons. As we previously reported, PANX1 protein localized primarily to the cell periphery of undifferentiated iPSCs, where it co-localized with actin (Figure 2A). PANX1 was similarly co-localized with actin in SOX2-expressing NPCs and TUJ1-expressing neurons (Figure 2A). NPCs in culture typically form polarized neural rosettes, identified by Nestin/SOX2 expression and characteristic flower petal arrangement (Wilson and Stice, 2006). Interestingly, as the NPCs in culture arranged into neural rosette-like structures, we observed PANX1 staining concentrated at the centermost (apical) side of the neural rosettes (Figure 2A). Western blotting revealed a significant upregulation of PANX1 protein as iPSCs differentiate toward NPCs and neurons (Figure 2B, C). Indeed, NPCs express 2.852 ± 0.522-fold more PANX1 and neurons express 5.324 ± 0.357-fold more PANX1 compared to undifferentiated iPSCs (Figure 2B, C). Additionally, we noted a difference in the PANX1 banding pattern on Western blots where NPCs and neurons possess a significantly greater proportion of the high molecular weight PANX1 isoform, most likely corresponding to the heavily glycosylated Gly2 species (Figure 2B, D). The putative Gly2 PANX1 species comprises 40.150 ± 0.843% of total PANX1 in iPSCs, 79.848 ± 1.551% in NPCs, and 84.370 ± 1.357% in neurons (Figure 2D). This dramatic upregulation of PANX1 protein during NPC and neuron differentiation suggests a role for PANX1 in neural specification and early human brain development.

**Figure 2.**
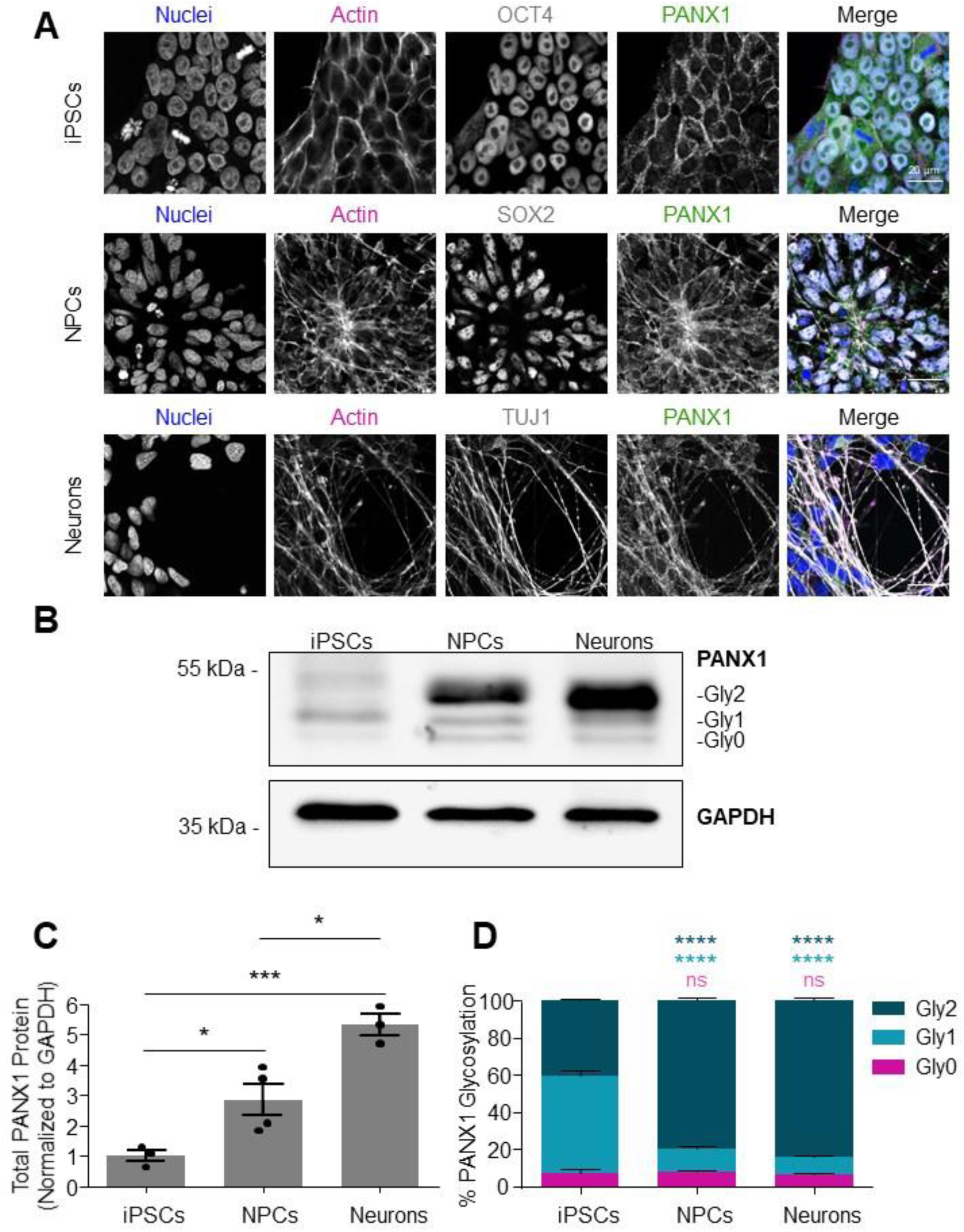
PANX1 protein expression is significantly upregulated following human iPSC differentiation to NPCs and neurons. **(A)** Representative immunofluorescence confocal micrographs depicting PANX1 protein (green) localization in OCT4-positive (gray) iPSCs, SOX2-positive (gray) NPCs, and TUJ1-positive (gray) neurons. Nuclei (Hoechst, blue); Actin (phalloidin, magenta). Scale bar = 20 μm. **(B)** Representative Western blot depicts PANX1 protein expression as three discrete bands corresponding to the putative heavily glycosylated (Gly2), high mannose (Gly1), and non-glycosylated species (Gly0) in human iPSCs, NPCs, and neurons. **(C)** Densitometry analysis of total PANX1 protein expression in iPSCs, differentiated NPCs and differentiated neurons. Data normalized to GAPDH and expressed as a fold of undifferentiated iPSCs. **(D)** Densitometric analysis of the proportion of Gly2, Gly1 and Gly0 PANX1 species expressed as a percent of the total PANX1 protein present. Error bars depict the standard error of the mean where data is representative of 3-4 independent experiments. *, p < 0.05; ***, p < 0.001; ****, p < 0.0001; ns = nonsignificant relative to iPSCs.

### PANX1 is apically expressed in budding neuroepithelia of iPSC-derived cerebral organoids

To further understand how PANX1 influences the earliest stages of human brain development, we next employed a cerebral organoid model to evaluate PANX1 localization throughout the embryonic and early fetal stages of human cortex development. Cerebral organoids are generated through 1) 3D induction of neuroectoderm from stem cell-derived embryoid bodies (EBs); 2) arrangement of neural rosettes and neuroepithelial expansion; 3) ventricular-like zone formation and intermediate progenitor emergence; 4) neuronal differentiation and cortical layering (Figure 3A). Ultimately, this results in a large, layered organoid comprised of numerous neural lineages.

**Figure 3.**
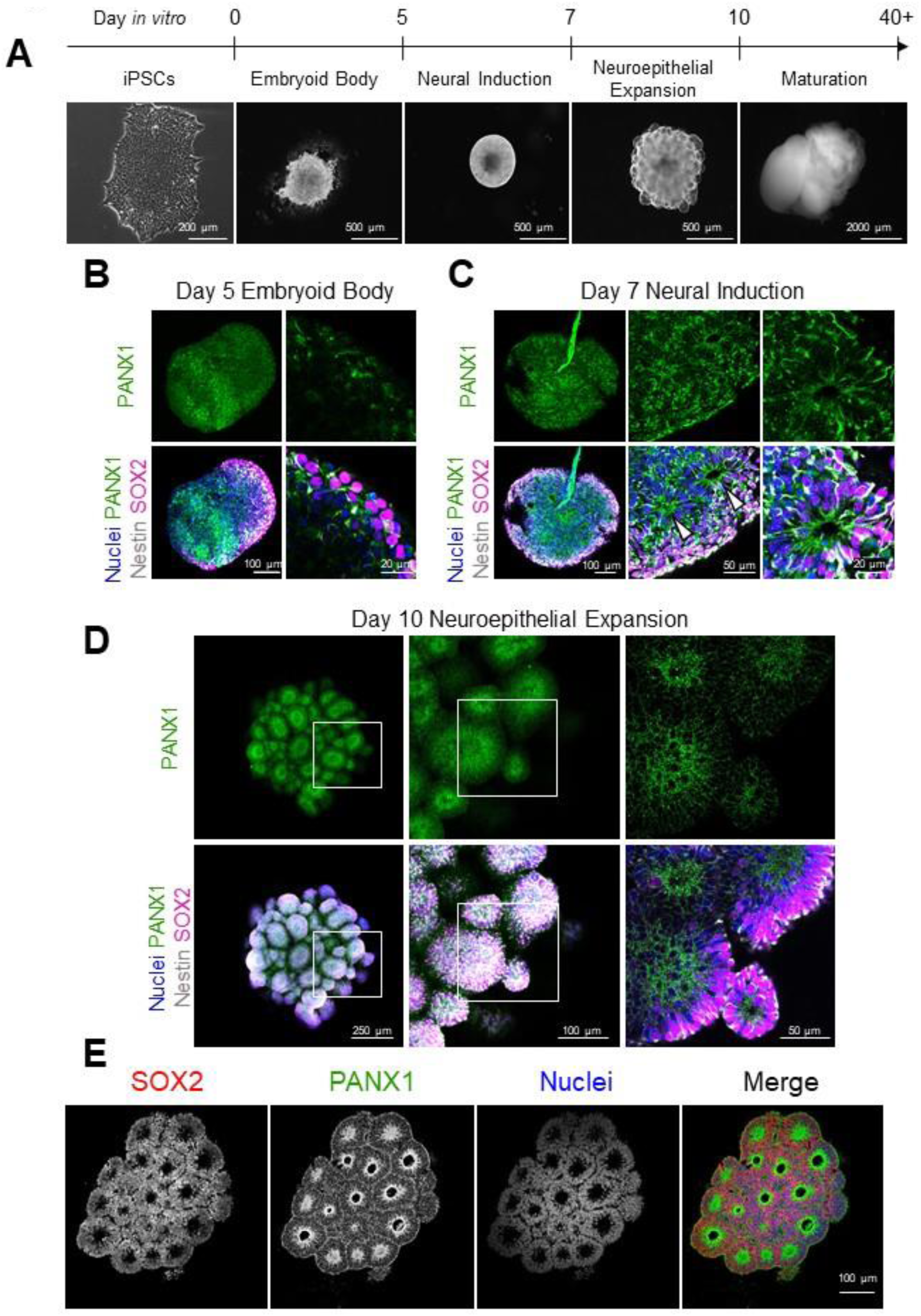
PANX1 is expressed at the earliest timepoints of cerebral organoid development. **(A)** Schematic depicting the various stages of cerebral organoid culture and representative phase images of organoid morphology. **(B-D)** Representative immunofluorescent confocal micrographs of whole mount control cerebral organoids at various developmental stages labelled for PANX1 (green), Nestin (gray), and SOX2 (magenta). **(B)** PANX1 is widely expressed across the EB stage of cerebral organoid development (day 5). **(C)** At the end of the neural induction stage (day 7), PANX1 is beginning to be apically localized in SOX2-positive neural rosettes (white arrowheads). **(D)** Day 10 cerebral organoids at the end of the neuroepithelial expansion stage develop many bulbous regions of radially arranged neuroepithelia with PANX1 staining concentrated at the center. **(E)** Immunofluorescence confocal imaging of cryosectioned day 10 cerebral organoids demonstrating radially arranged SOX2 positive neuroepithelia (red) with PANX1 staining (green) concentrated at the center. Scale bars as indicated. Nuclei (Hoechst, blue).

At the end of the EB stage (day 5), organoids appear as a dense, disorganized cellular mass exhibiting some SOX2 and Nestin-positive regions. Whole mount immunofluorescence confocal microscopy revealed wide PANX1 expression throughout the EB (Figure 3B). Upon the initiation of neural induction (day 7), organoids begin to arrange into pseudostratified neural rosettes lined by polarized SOX2-positive cells. At this stage, the organoid consists of multiple rosette-like arrangements of SOX2-positive neuroepithelial cells surrounding fluid-filled spaces. Similar to what we observed in the 2D NPC cultures, PANX1 staining shifted at this stage to concentrate at the apical membrane region of these neural rosettes (Figure 3C, arrow heads). As apical-basal polarity becomes fully established at the end of neuroepithelial expansion (day 10), PANX1 is preferentially localized toward the apical surface of each neuroepithelial sphere as shown through whole mount immunofluorescence (Figure 3D). To complement the whole mount immunofluorescence data presented in Figure 3B-D, we cryosectioned day 10 organoids and performed immunofluorescence confocal imaging of PANX1 at the neuroepithelial expansion stage of cortical organoid development. Confocal imaging of cryosectioned day 10 organoids confirmed a striking concentration of PANX1 protein localized to the apical edge of the neuroepithelial buds (Figure 3E). Together, these data indicate that PANX1 is expressed throughout the embryonic stages of cerebral organoid development, from the beginning of neural lineage commitment to the formation of polarized neuroepithelium where the channels largely reside at the apical edge.

### PANX1 genetic ablation and pharmacological inhibition results in significantly smaller organoids

At neuroepithelial expansion, NPCs proliferate rapidly via symmetric division to make up the required tissue and organ mass. After expansion, NPCs must successfully migrate and differentiate via asymmetric division into mature neural cell types. We have previously reported that *PANX1-/-* iPSCs exhibit deficits in ectoderm lineage specification (Noort et al., 2021). Given the upregulation of PANX1 during NPC differentiation, we hypothesized that loss of PANX1 would compromise the neuroepithelial expansion stage of organoid development. Immunofluorescence imaging of whole mount day 10 organoids confirmed the absence of PANX1 in our CRISPR-Cas9 knockout organoids (Figure 4A). *PANX1-/-* organoids were significantly smaller than control, which was even more pronounced in probenecid (PBN)-treated organoids (Figure 4B). Possible reasons for smaller organoids at neuroepithelial expansion include differences in apoptosis, proliferation, or an imbalance in symmetric/asymmetric cell division. We found similar proportions of cleaved caspase 3 and ki67 positive cells within control and *PANX1-/-* organoids, suggesting little difference in apoptosis or cell proliferation (Figure 4C). Cell division angle relative to the apical surface can reveal whether NPCs will undergo symmetrical division (self-renewal) or asymmetrical division (differentiation) (Wilson and Stice, 2006). Because PANX1 is reported to positively regulate neural progenitor cell self-renewal and proliferation (Wicki-Stordeur et al., 2012), we examined organoid size and thickness of the PAX6+ neuroepithelial progenitors in day 10 organoids. Skewed symmetrical/asymmetrical NPC division could result in premature neuronal differentiation, which could account for the small size of *PANX1-/-* organoids. However, we found no difference in the proportion of nuclei undergoing vertical (symmetrical) or horizontal (asymmetrical) divisions (Figure 4D-F). Furthermore, the thickness of the PAX6+ progenitor cell layer was not significantly different in *PANX1-/-* organoids compared to control (Figure 4G).

**Figure 4.**
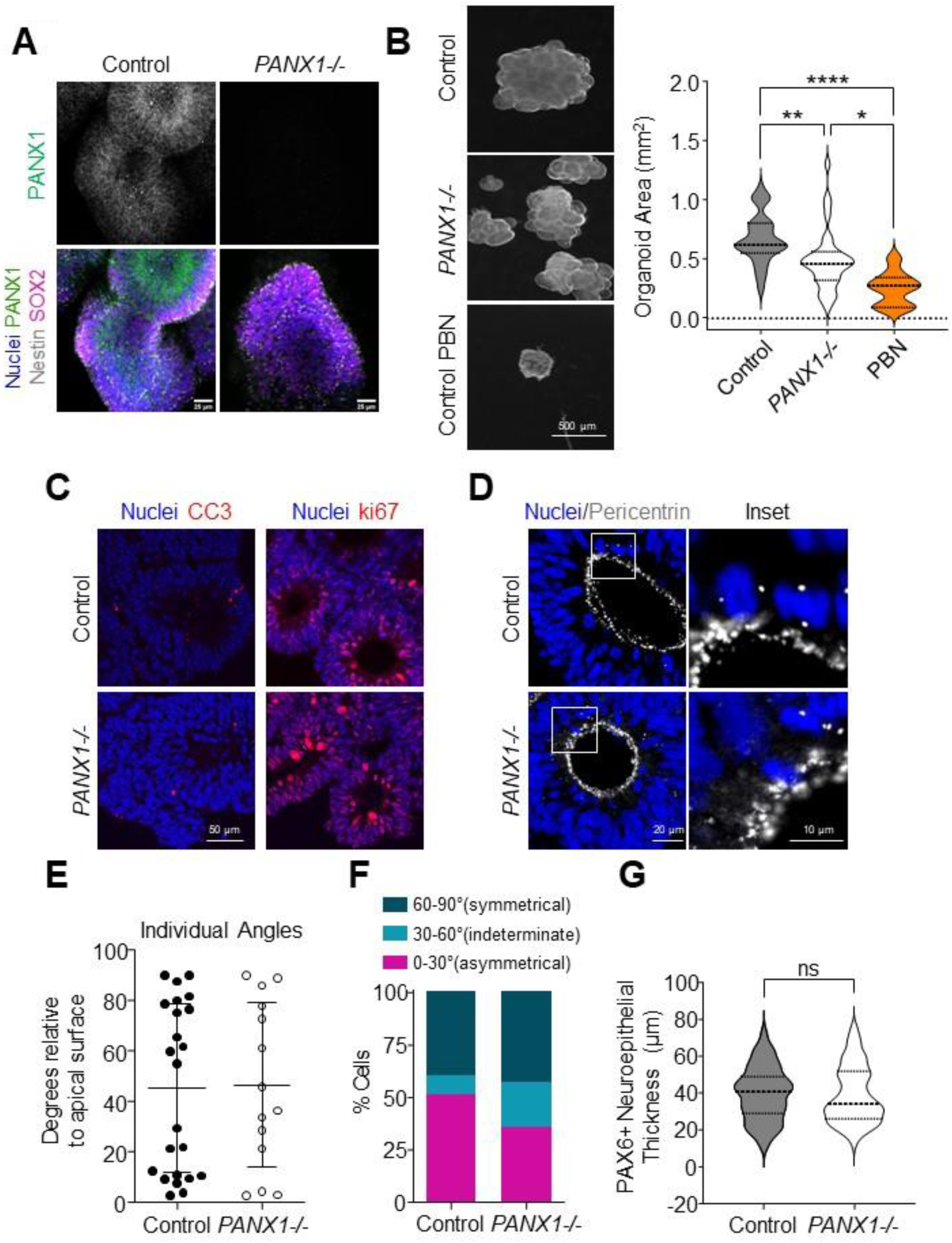
PANX1 inhibition results in significantly smaller organoids at neuroepithelial expansion stage. **(A)** Whole mount immunofluorescence confocal imaging of PANX1 (green) in control and *PANX1-/-* day 10 cerebral organoids. Nestin (gray), SOX2 (magenta), nuclei (Hoechst, blue). **(B)** Representative phase images and analysis of organoid area of control and *PANX1-/-* organoids at day 10 as well as control organoids treated with 1 mM probenecid (PBN) starting on day 5. N=46 for untreated control organoids, N=34 for untreated *PANX1-/-* organoids, N=9 for control probenecid-treated organoids. *, p < 0.05; **, p < 0.01; ****, p < 0.0001. **(C)** Representative immunofluorescence confocal micrographs of cleaved caspase 3 (CC3) or ki67 (red) and nuclei (Hoechst, blue) in day 10 control and *PANX1-/-* organoids. **(D)** Representative immunofluorescence confocal micrographs of pericentrin (gray) and nuclei (Hoechst, blue) in day 10 control and *PANX1-/-* organoids. Pericentrin reveals the angle of division in neuroepithelial cells at the apical domain of control and knockout organoids. **(E-F)** Categorization of cell division angles relative to the apical surface. Angles of 0-30’ indicate horizontal cleavage (asymmetric divisions and differentiation of the daughter cell). Angles of 30-60 degrees are indeterminate. Angles of 60-90’ indicate vertical cleavages (symmetrical divisions resulting in NPC renewal). **(G)** Thickness of PAX6+ neuroepithelial cells in control and *PANX1-/-* day 10 organoids (from apical to basolateral edge). Control = 81 FOV; *PANX1-/-* = 84 FOV. Scale bars as indicated.

### Transcriptomic analysis of PANX1-/- organoids reveal gene expression changes related to neural development

Although PANX1 genetic ablation and pharmacological inhibition resulted in significantly smaller organoids at the neuroepithelial stage, none of the metrics we evaluated in Figure 4 were changed in our *PANX1-/-* organoids. Therefore, we compared transcriptomic profiles of day 10 *PANX1-/-* organoids to control organoids using RNA-sequencing technology. Immunofluorescence imaging of cryosectioned day 10 organoids confirmed that nearly all of cells within day 10 cerebral organoids are PAX6 positive neural progenitor cells (Figure 5A). Analysis of transcriptomic data using DESeq2 revealed a total of 1,047 differentially expressed genes with an adjusted p-value of less than 0.05. Limiting the genes to those with a log fold change (log_2_FC) of at least 1 (or −1) resulted in 453 differentially expressed genes (231 up-regulated, 222 down-regulated) in response to *PANX1* knockout (Figure 5B; Supplementary Data). Similar to our previous study, pluripotency-related genes such as *POU5F1* (OCT4) and *ZSCAN10* were among the most significantly upregulated individual genes in *PANX1-/-* organoids compared to control (Noort et al., 2021). Also observed was upregulation of vertebrae development-associated (*VRTN*), cell differentiation homeobox protein (*NKX1-2*) (Brandenberger et al., 2004), and developmental factor Forkhead box H1 (*FOXH1*) (Pluta et al., 2022). The most significantly downregulated genes included the anti-apoptotic coiled-coil-helix-coiled-coil-helix domain containing 2 (*CHCHD2*) (Zhu et al., 2016), the orphan nuclear receptor tailless (*TLX/NR2E1*) (Islam and Zhang, 2015), signaling molecule R-spondin 2 gene (*RSPO2*) (Gyllborg et al., 2018) and neuronal homeobox genes *BARHL1* and *BARHL2* (Li et al., 2004).

**Figure 5.**
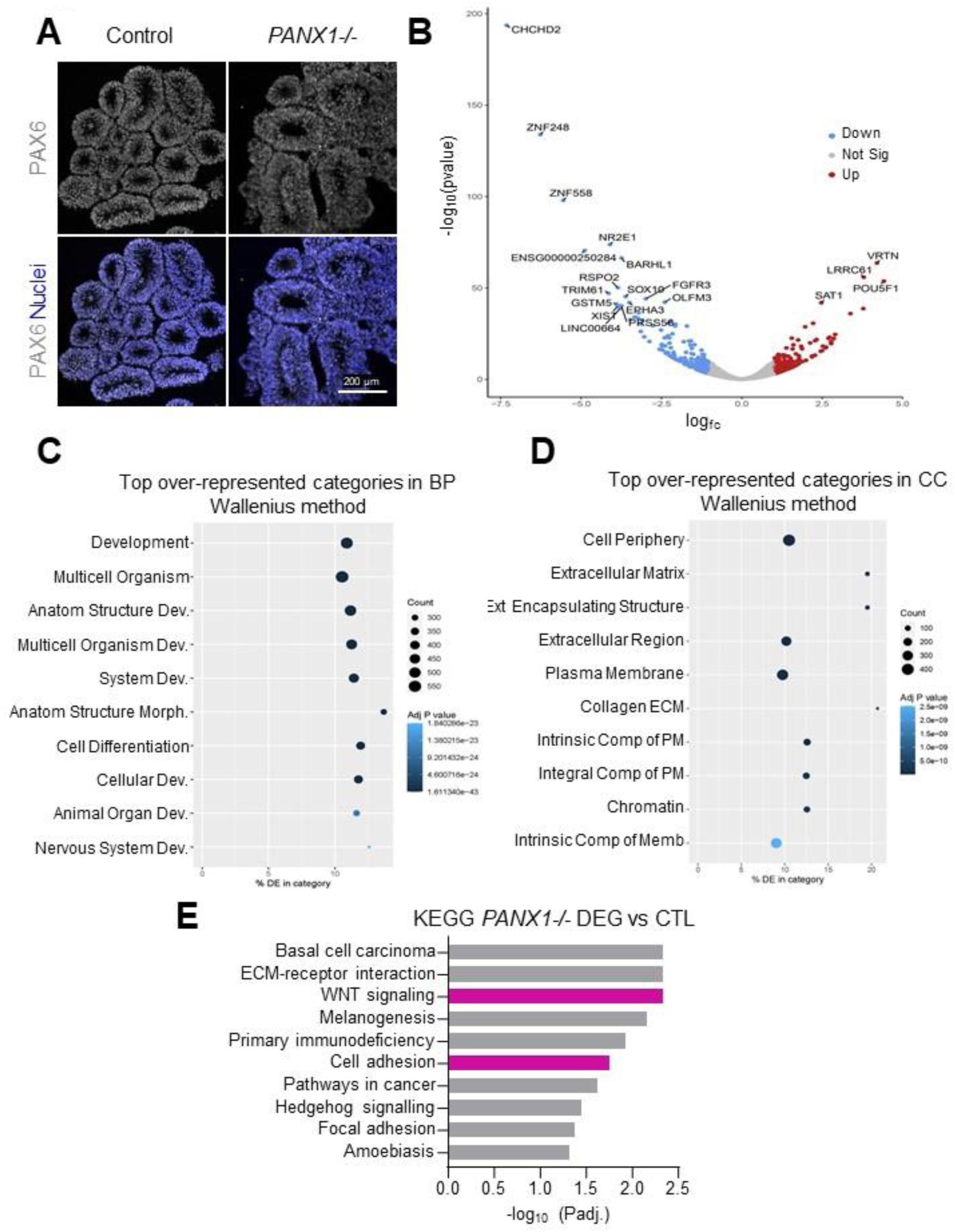
*PANX1-/-* organoids exhibit gene expression changes related to cell-cell adhesion, signaling, extracellular matrix and development. **(A)** Immunofluorescence confocal imaging of cryosectioned day 10 cerebral organoids demonstrating proportion of PAX6 positive (gray) neural progenitor cells. Scale bars as indicated. **(B-E)** Bulk RNA sequencing of control and *PANX1-/-* day 10 cerebral organoids. **(B)** Volcano plot illustrating the top differentially expressed genes where log2 fold change is >|1|. 231 genes were significantly UP-regulated in *PANX1-/-* relative to control and 222 genes were significantly DOWN-regulated. **(C)** GoSeq plot showing the top over-represented categories in biological process. **(D)** GoSeq plot showing the top over-represented categories in cellular compartment. **(E)** KEGG pathway analysis for significant differentially expressed genes in *PANX1-/-* organoids. Highlighted in pink are KEGG pathways associated with WNT signaling and cell adhesion.

Gene set enrichment analysis (GSEA) using gene ontology (GO) terms related to biological processes (BP), molecular function (MF), cellular component (CC), and KEGG pathways was used to group differentially expressed genes along common biological themes (Figure 5C-E; Supplementary Data). Most GO:BP categories included genes involved in developmental processes (Figure 5C) while GO:MF categories were over-represented by genes associated with cell signaling. In terms of cellular components, differentially expressed genes were over-represented in GO categories related to plasma membrane components as well as extracellular and peripheral cellular regions (Figure 5D). GSEA analysis allowed us to categorize differentially expressed genes according to KEGG pathway maps representing what is currently known about molecular interactions and biological networks. We identified 10 KEGG pathways that were significantly over-represented in our GSEA analysis (adjusted p-value <0.05) (Figure 5E and Supplementary Data). Among the most over-represented KEGG pathways are those associated with ECM-receptor interactions and adhesion molecules (KEGG categories 04512, 04514 and 04510) (Table 2). Signaling pathways related to WNT signaling and Hedgehog signaling were also represented in our differentially expressed genes (KEGG categories 04310 and 04340) (Table 3). Finally, we found a surprising number of non-coding RNA molecules differentially expressed in our *PANX1-/-* organoids compared to control (Table 4).

**Table 2.** Differentially expressed adhesion genes in day 10 *PANX1-/-* organoids.

| Adhesion Up | Fold increase | Role in neurodevelopment | Refs |
| --- | --- | --- | --- |
| *PCDHGB4* | 3.011 | Pathogenic sequence variations related to neurodevelopmental disorder with poor growth and skeletal anomalies<br>Involved self-recognition and avoidance | (Neel et al., 2022) |
| *CD-40* | 2.603 | Hippocampal excitatory neuronal dendrites stunted in *Cd40*<sup>-/-</sup> mice<br>CD40 isoforms temporally modulate neuron differentiation during brain development | (Hou et al., 2008; Carriba and Davies, 2017) |
| *CNTN2* | 2.392 | Axonal elongation, axonal guidance, and cellular migration<br>*CNTN2*/*TAG-1* deficient mice exhibit a significantly smaller cortex and a reduction of corticothalamic axons | (Masuda, 2017; Kastriti et al., 2019) |
| *PCDHGA10* | 2.356 | *Pcdhg* isoforms regulate cortical inhibitory interneuron survival during the endogenous period of programmed cell death | (Carriere et al., 2020; Mancía Leon et al., 2020) |
| *CLDN6* | 2.229 |  |  |
| *PCDH8* | 2.204 | *Pcdh8* knockout mouse neurons exhibit increased dendritic spines in the presence of N-cadherin | (Yasuda et al., 2007; Takeuchi et al., 2020) |
| *CD4* | 2.015 |  |  |
| *PCDHA10* | 2.007 | *Pcdhg* isoforms regulate cortical inhibitory interneuron survival during the endogenous period of programmed cell death | (Carriere et al., 2020; Mancía Leon et al., 2020) |

| Adhesion Down | Fold decrease | Role in neurodevelopment | Refs |
| --- | --- | --- | --- |
| *PCDH7* | 2.557 | Promotes neural differentiation and dendritic spine morphology in zebrafish and mouse development | (Wang et al., 2020b; Xiao et al., 2023) |
| *CDH15* | 2.353 | Linked to ASD and Intellectual Disability | (Bhalla et al., 2008; Zarrei et al., 2019) |
| *VCAM1* | 2.128 | Present in adult hippocampal neural stem cell subpopulation<br>*Vcam1*<sup>-/-</sup> mice exhibit impaired spatial learning and memory in mice and fewer proliferating NSCs | (Wang et al., 2020a) |
| *CDH9* | 2.108 | GWAS association with ASD | (Wang et al., 2019) |
| *CDH19* | 2.047 |  |  |
| *NCAM2* | 2.015 | Implicated in neurodevelopmental disorders including Down syndrome and autism<br>Promotes filopodia formation and neurite branching<br>*NCAM2* downregulation leads to dendritic branching defects | (Sheng et al., 2015; Parcerisas et al., 2020) |
| *CLDN1* | 1.805 | Patients with *CLDN1* mutations may be an increased risk for learning disabilities, mental retardation, and language delay | (Salik et al., 2022) |
| *PCDHA11* | 1.733 | Pcdhg isoforms regulate cortical inhibitory interneuron survival during the endogenous period of programmed cell death | (Carriere et al., 2020; Mancía Leon et al., 2020) |
| *MPZ* | 1.331 | Adhesion molecule necessary for normal myelination in the peripheral nervous system<br>Associated with Charcot-Marie-Tooth neuropathy type 1 B (CMT1B) | (Speevak and Farrell, 2013; Sanmaneechai et al., 2015) |

**Table 3.** Differentially expressed WNT pathway genes in day 10 *PANX1-/-* organoids.

| WNT up | Fold increase | Role in neurodevelopment | Refs |
| --- | --- | --- | --- |
| *WNT8A* | 5.49 | Wnt8 inhibits BMP4 expression, allowing neural induction from ectoderm<br>Promotes synaptogenesis via LRP6 | (Baker et al., 1999; Sharma et al., 2013) |
| *DKK4* | 3.61 | Associated with schizophrenia | Proitsi et al., 2008 |
| *FZD10* | 2.52 | Plays a role in neural tube patterning | (Alrefaei et al., 2020) |
| *PRKG2* | 2.49 | PRKG2-deficient mice exhibit restricted growth and deficits in learning and memory, similar to human patients | (Tran et al., 2021) |
| *RSPO4* | 2.23 |  |  |

| WNT down | Fold decrease | Role in neurodevelopment | Refs |
| --- | --- | --- | --- |
| *RSPO2* | 14.47 | Promotes Midbrain Dopaminergic Neurogenesis and Differentiation | (Gyllborg et al., 2018) |
| *WNT2B* | 4.91 | Associated with bipolar disorder | (Zandi et al., 2008) |
| *WNT7A* | 4.72 | Stimulates neural stem cell proliferation and promotes neuronal differentiation<br>Associated with bipolar disorder | (Zandi et al., 2008; Qu et al., 2010; Qu et al., 2013) |
| *RSPO3* | 4.22 |  |  |
| *RSPO1* | 2.99 |  |  |
| *SOST* | 2.34 |  |  |
| *WNT7B* | 2.26 | Promotes dendrite development in mouse hippocampal neurons | (Rosso et al., 2005; Ferrari et al., 2018) |
| *WNT8B* | 2.15 | Expression restricted to the developing forebrain | (Lako et al., 1998) |
| *WNT3A* | 2.01 |  |  |

**Table 4.** Differentially expressed non-coding RNAs in day 10 *PANX1-/-* organoids.

| ncRNA Up | Fold increase | Role in neurodevelopment | Refs |
| --- | --- | --- | --- |
| *JAKMIP2-AS1* | 5.74 | High expression in neural precursor cells | (Yan et al., 2023) |
| *MIR302CHG* | 3.46 | MiR-302/367 cluster host regulates neural progenitor proliferation, differentiation and survival during neurulation | (Yang et al., 2015) |
| *PCAT14* | 3.16 |  |  |
| *LINC01356* | 3.00 |  |  |
| *LINC00958* | 2.41 |  |  |
| *LINC00649* | 2.39 |  |  |
| *MIR100HG* | 2.27 |  |  |
| *LINC02381* | 2.15 |  |  |
| *LNCTAM34A* | 2.11 |  |  |
| *SATB1-AS1* | 2.10 | Antisense for SATB1 (TF associated with neuronal maturation and cortical development)<br>Associated with Anxiety disorder | (Denaxa et al., 2012; Levey et al., 2020) |
| *XACT* | 2.06 | Regulates neuronal differentiation in both male and female hPSCs | (Motosugi et al., 2021) |
| *LINC01415* | 2.01 |  |  |
| *LINC01833* | 2.01 |  |  |

| ncRNA Down | Fold decrease | Role in neurodevelopment | Refs |
| --- | --- | --- | --- |
| *XIST* | 14.49 | XIST dosage correction of Trisomy 21 promotes NSC differentiation to neurons | (Czerwiński and Lawrence, 2020) |
| *LINC00664* | 13.77 |  |  |
| *LINC01515* | 5.62 | Dysregulated in autism spectrum disorder patients | (Mordaunt et al., 2020) |
| *SVIL-AS1* | 5.50 |  |  |
| *MIR9-1HG* | 4.63 | Potential regulator of primate ganglionic eminences | (Zhao et al., 2022) |
| *FAM218A* | 4.54 |  |  |
| *FLG-AS1* | 4.21 |  |  |
| *MIR9-3HG* | 2.63 |  |  |
| *LINC00461* | 2.53 | Pleiotropic effects on five psychiatric traits; important for mouse neurodevelopment | (Liu et al., 2020) |
| *OSTM1-AS1* | 2.50 |  |  |
| *MIR1-1HG-AS1* | 2.44 |  |  |
| *MIR924HG* | 2.43 |  |  |
| *NR2F1-AS1* | 2.30 | Pro-neurogenic; genetic disruption contributes to complex neurodevelopmental disorders. | (Ang et al., 2019) |
| LHX5-AS1 | 2.06 | Strongly enriched in early human embryonic brain, becoming restricted to the developing cortical plate in stage 14-16 embryos | (Eze et al., 2021) |

### PANX1 co-localizes with apically-situated junctional proteins at the neuroepithelial expansion stage of cerebral organoids

Given the abundance of altered genes related to cell adhesion and WNT signaling in our *PANX1-/-* organoids, and because β-catenin was recently recognized as a PANX1 interacting partner (Sayedyahossein et al., 2021), we investigated key junctional protein targets at the apical domain to see if they are differentially expressed or localized in *PANX1-/-* organoids (Figures 6, 7). Rosette formation depends upon apical-basolateral patterning, coordinated by several key proteins including N-Cadherin, β-Catenin and ZO-1 (Wilson and Stice, 2006). Therefore, we next examined the co-localization of PANX1 with these apically-situated adhesion proteins (Figure 6). Manders’ correlation coefficients demonstrated a robust colocalization of PANX1 with β-catenin (0.4070 ± 0.06509) and PANX1 with N-cadherin (0.5457 ± 0.05012). However, there was very little colocalization between PANX1 and the tight junction protein ZO-1 (0.06847 ± 0.01272) (Figure 6C). We next evaluated the expression and localization of β-catenin, Claudin1, N-cadherin and ZO-1 in *PANX1-/-* organoids (Figure 7). Although several of these gene classes were significantly altered in the RNAseq data set, Western blotting and immunofluorescence showed similar expression and localization patterns between control and *PANX1-/-* organoids (Figure 7A-C). Claudin 1 localization was a bit trickier as it was only observed at the apical membrane domain in ∼33% of control neural rosettes (Figure 7A, B, D). However, Claudin 1 was never present at the apical membrane domain in any *PANX1-/-* neural rosettes.

**Figure 6.**
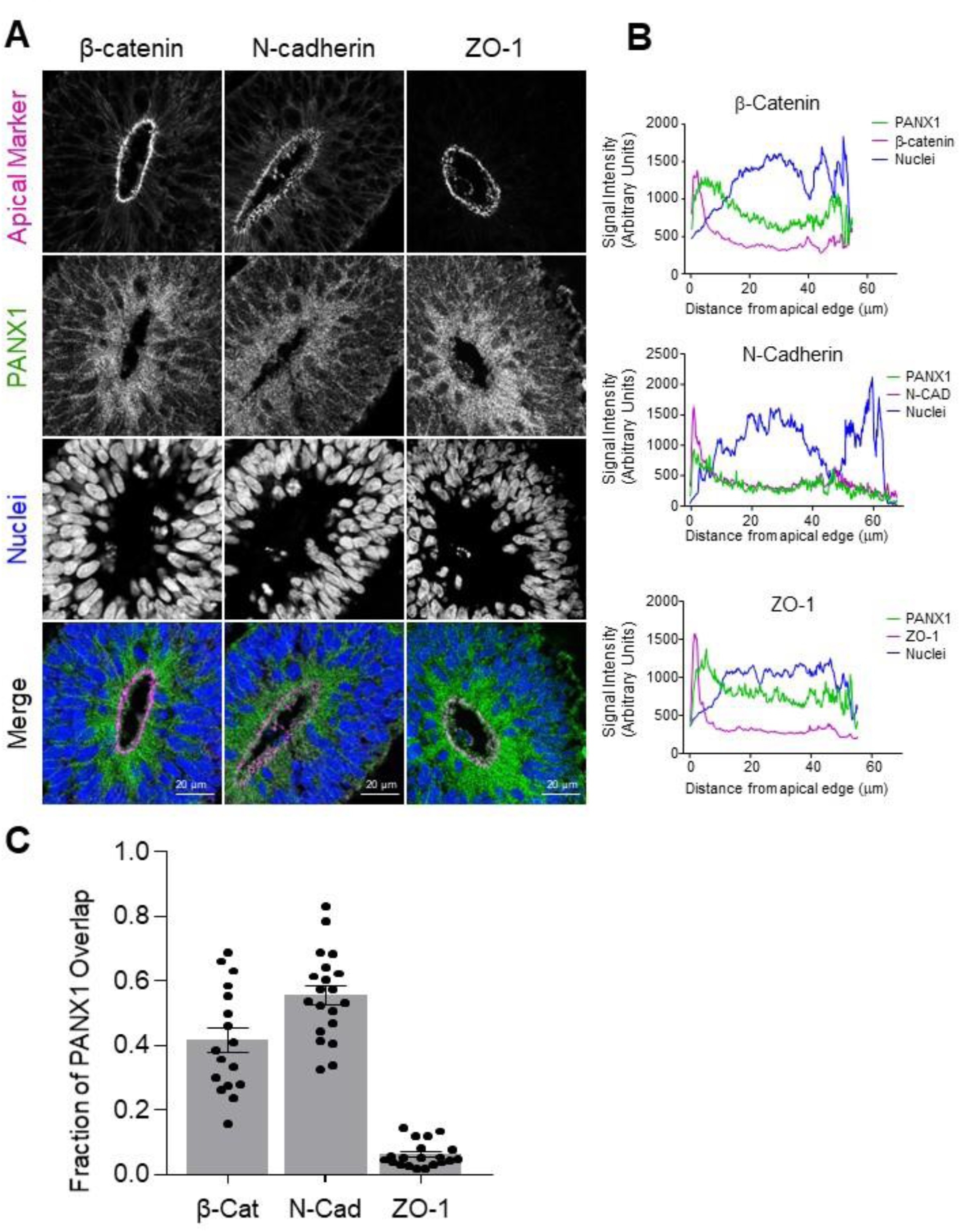
PANX1 co-localizes with key junctional proteins at the apical edge of neuroepithelial cells in cerebral organoids. **(A)** Representative immunofluorescence confocal images demonstrating PANX1 (green) co-localization with the apical membrane proteins β-catenin, N-cadherin, and ZO-1 (magenta) at the apical side (innermost) of the neuroepithelia. Scale bars as indicated. **(B)** Line graphs and **(C)** Manders’ correlation coefficients depicting the fraction of signal overlap for PANX1 with apically situated adherens junctions β-catenin and N-cadherin as well as with tight junction ZO-1. Error bars depict the standard error of the mean for 17-24 fields of view obtained from 4 independent experiments.

**Figure 7.**
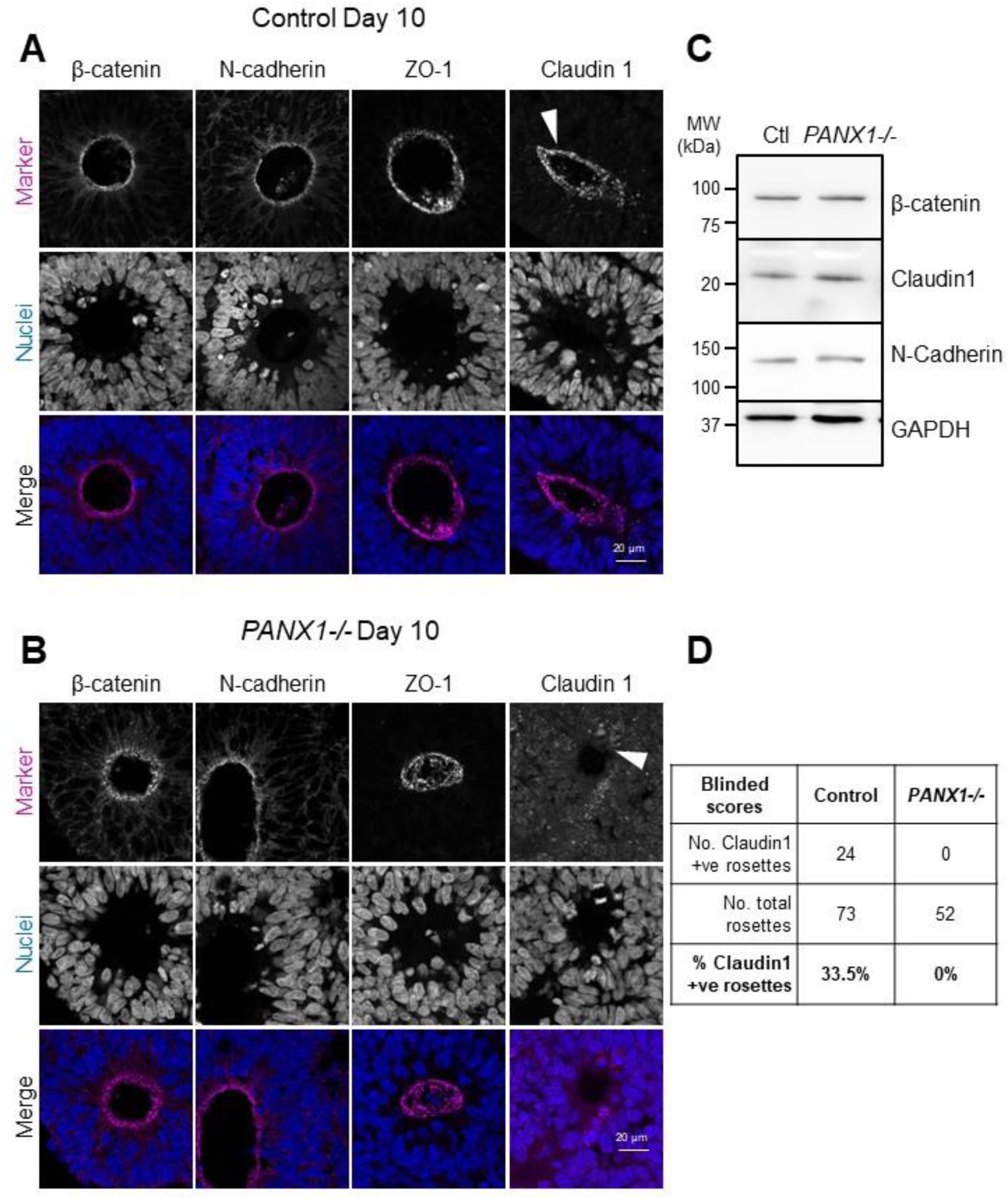
Apical adhesion molecule expression and localization unchanged in *PANX1-/-* cerebral organoids. **(A, B)** Representative immunofluorescence confocal micrographs for select apical markers (magenta) in day 10 control and *PANX1-/-* cerebral organoids. Nuclei (Hoechst, blue). Arrowhead indicates the re-distribution of Claudin 1 away from the apical membrane domain in *PANX1-/-* organoids. Scale bars as indicated. **(C)** Western blots of apical protein expression in control and *PANX1-/-* day 10 cerebral organoids. **(D)** Blinded scoring of Claudin 1 localization to apical membrane domain in control and *PANX1-/-* day 10 cerebral organoids.

### PANX1 expression is preferentially expressed in neurons as cerebral organoids mature

As cerebral organoids mature, the numerous neural rosettes continue to hollow out and elongate, forming fluid-filled ventricular-like spaces. Furthermore, the tightly packed neuroepithelia and NPCs residing in the emerging ventricular-like zone begin to asymmetrically divide and migrate, ultimately differentiating into neurons, and later, to glia. To examine how PANX1 expression and localization change as organoids begin to mature and establish cortical layering, we examined mature cerebral organoids between 40 to 120 days (Figure 8). Immunostaining revealed some apical expression of PANX1 along SOX2-positive ventricular-like zones in day 40 organoids (Figure 8A). However, PANX1 signal intensity appeared brightest outside the ventricular-like zones coinciding in regions with TUJ1-positive neurons (Figure 8B). In 120-day old organoids we observed PANX1 expression within stellate GFAP-positive cells with astrocyte-like morphology (Figure 8C). Our results indicate that mature organoids exhibit PANX1 localization at the apical side of the ventricular-like zones and in more developmentally advanced neural cell types such as neurons and GFAP-positive glia.

**Figure 8.**
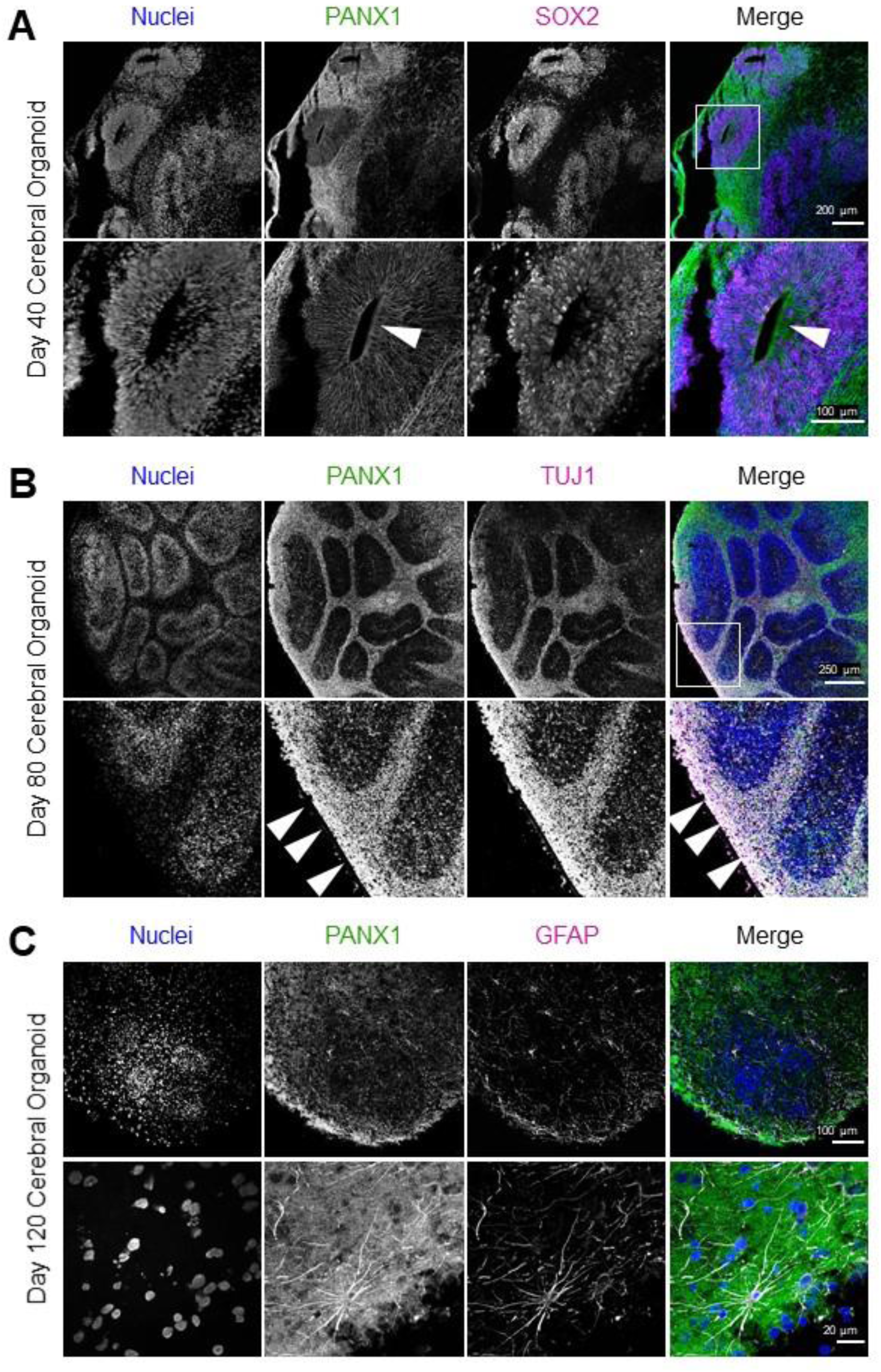
PANX1 expression in neurons and glia of mature cerebral organoids. Representative immunofluorescence confocal images showing PANX1 (green) localization to various regions and cell types throughout cerebral organoid maturation**. (A)** Forty-day-old cerebral organoids exhibit abundant PANX1 staining outside the SOX2-positive ventricular-like zones with limited PANX1 expression persisting at the ventricular-like zone’s apical edge (arrowhead, inset). **(B)** Day 80 cerebral organoids display bright PANX1 expression in areas where TUJ1-positive neurons reside (arrowheads, inset). **(C)** PANX1 expression is apparent in GFAP-positive glial cells with astrocyte-like morphology in 120-day-old cerebral organoids. Nuclei (Hoechst, blue). Neural cell type markers (SOX2, TUJ1, and GFAP in magenta). Scale bars as indicated.

## Discussion

Given that PANX1 is expressed in the earliest cell types of human development and is linked to neurological disease, we sought to explore PANX1 expression and localization throughout early stages of human brain development. Although most PANX1 studies focus on perinatal or adult mouse models, PANX1 is expressed in human oocytes, embryos and pluripotent stem cells, suggesting a fundamental role for PANX1 in human development (Hainz et al., 2018; Sang et al., 2019; Noort et al., 2021). The potential role of PANX1 in human brain development is further supported by a loss-of-function human germline *PANX1* variant in a patient with severe neurological deficits (Shao et al., 2016), and by *PANX1* and *PANX2* single nucleotide polymorphisms implicated in autism spectrum disorder (Davis et al., 2012). Indeed, we recently reported that PANX1 is expressed in human induced pluripotent stem cells (iPSCs) and our *PANX1-/-* iPSCs exhibit decreased ectoderm formation compared to control (Noort et al., 2021). This finding indicated to us that PANX1 might impact the development of ectodermal-derived tissues, such as the brain. The Allen Institute’s Brainspan prenatal laser microdissection microarray dataset depicts PANX1 transcript expression in 21 pcw human fetal brains (Brainspan.org). Because transcript expression does not always correlate to protein expression, we were surprised that we could not find published evidence of PANX1 protein expression at this stage of human development. We have now confirmed that PANX1 protein is also expressed across all layers of the developing human cerebral cortex with brighter manifestation in the marginal zone and subventricular zone. The high PANX1 expression in the human fetal brain, combined with our findings of ectodermal lineage deficits in human *PANX1-/-* iPSCs suggests a role for PANX1 in human neural development.

We previously reported that PANX1 is expressed at the cell surface across iPSCs and iPSC-derived embryoid bodies (Noort et al., 2021). Intriguingly, we find here that PANX1 localization becomes very restricted upon cerebral organoid neural induction, when organoids begin to arrange into neural rosettes (Figures 2, 3, 6). As apical-basolateral polarity becomes fully established at the end of neuroepithelial expansion, the organoid consists of multiple rosette-like arrangements of SOX2-positive neuroepithelial cells surrounding fluid-filled spaces. These neuroepithelia are the progenitor cells of the developing brain and express tight junctions, adherens junctions, and exhibit apical-basolateral polarity, forming a layer of pseudostratified columnar neuroepithelium which gives rise to the neural plate and subsequent neural tube (Mori et al., 2005; Wilson and Stice, 2006; Kriegstein and Alvarez-Buylla, 2009). At the neuroepithelial expansion stage of cerebral organoid development, PANX1 was preferentially localized toward the apical surface of each neuroepithelial rosette and colocalized with key apical proteins including β-catenin and N-cadherin (Figure 6). Our RNAseq in day 10 organoids revealed a significant downregulation in several adhesion molecules including cadherins, claudins, neural cell adhesion molecule 1 (NCAM), and others. In addition to establishing polarity in the developing brain, these cytoskeleton-anchoring proteins help to coordinate the mitotic spindle orientation and several downstream signal transduction cascades controlling neural cell fate. In the future it would be interesting to investigate whether PANX1 physically interacts with these junctional proteins as has been recently shown in other systems (Sayedyahossein et al., 2021). In addition to being significantly upregulated in neurons compared to iPSCs, our Western blot analyses showed differential PANX1 banding patterns between iPSCs, NPCs, and neurons where most of the PANX1 in NPCs and neurons exists as a high molecular weight species. These different molecular weight species have been shown to correspond to different PANX1 glycosylation species with unglycosylated (Gly0), simple glycosylation (Gly1) and complex carbohydrate modifications (Gly2), which are thought to influence plasma membrane targeting (Penuela et al., 2009). Other groups have shown that PANX1 may be preferentially distributed to basolateral or apical membrane compartments depending on the cell and tissue type where intracellular PANX1 retention inhibits cell polarization (Turmel et al., 2011; Shum et al., 2019). Apical PANX1 channels residing at the unopposed edge of the ventricular-like zone may have implications for paracrine signaling and long-range coordination of NPC proliferation in the ventricular-like zone. In the postnatal murine brain, ATP released into the extracellular space by PANX1 channels activates P2X7 and P2Y purinergic receptors which in turn stimulate the proliferation of NPCs (Wicki-Stordeur et al., 2012). Future studies will determine whether PANX1 serves a similar role in neuroepithelial expansion at this early stage of development.

Current literature implicates PANX1 with Wnt/β-catenin signaling through the physical interaction of PANX1 with β-catenin in melanoma cells (Sayedyahossein et al., 2021). Here we find that PANX1 and β-catenin are both apically localized in day 10 neuroepithelial rosettes and exhibit considerable co-localization. In the brain, Wnt signaling inhibits the self-renewal of cortical neural precursor cells and promotes differentiation of neuronal cell types such as dopaminergic neurons (Hirabayashi et al., 2004; Gyllborg et al., 2018). Indeed, several of the differentially expressed Wnt-associated genes in our *PANX1-/-* organoids are associated with neurodevelopment and neural stem cells.

Our RNAseq screen revealed a surprising number of differentially expressed non-coding RNAs (ncRNAs). Very few studies to date have linked PANX1 with ncRNA expression (Kim et al., 2021; Montagne et al., 2022). Not all of the ncRNAs that came out in our screen have documented roles in neurodevelopment, however several are associated with neural stem cells, neuronal differentiation and neurodevelopmental disorders. Many of the ncRNAs in our screen are related to cell cycle control and apoptosis in different forms of cancer. Given the smaller size of our *PANX1-/-* day 10 organoids, it is possible that differential expression of ncRNAs could contribute to changes in neural precursor cell proliferation or apoptosis. However, as we saw no obvious changes in ki67 or cleaved caspase 3 expression in our *PANX1-/-* organoids, we suspect the smaller organoids are not related to changes in cell proliferation or apoptosis.

Despite being apically localized at the neuroepithelial expansion stage, PANX1 resided primarily in TUJ1-expressing neurons in mature cerebral organoids, with lesser amounts persisting in the SOX2-positive ventricular-like regions (Figure 8). Consistent with this observation, PANX1 protein expression was significantly elevated in differentiated neurons compared to NPCs (Figure 2). We found a similar pattern in the human fetal cortex where PANX1 staining was concentrated outside of the ventricular zone (Figure 1). This is in contrast to reports in mouse brains where PANX1 was found to be concentrated in periventricular neural stem cells in postnatal day 15-60 mice (Wicki-Stordeur et al., 2012; Wicki-Stordeur and Swayne, 2013; Wicki-Stordeur et al., 2016). Just as we found a dramatic shift in PANX1 cellular distribution between day 10 and 40 cerebral organoids, it is possible that PANX1 could again change distribution between the fetal and adult brains.

In postnatal murine brains, pharmacological inhibition of PANX1 channels prevents NPC proliferation and enhances neuronal differentiation by promoting neurite extension and cell migration (Wicki-Stordeur et al., 2012; Wicki-Stordeur and Swayne, 2013). Others have demonstrated PANX1 localization at neuronal synapses where the channels help to replenish extracellular ATP, negatively regulate dendritic spine density, and maintain synaptic strength (Prochnow et al., 2012; Sanchez-Arias et al., 2020). (Penuela et al., 2014). It remains to be seen whether human iPSC-derived *PANX1-/-* neurons exhibit similar increases in spine and branching density as has been observed in mouse.

A caveat of human cerebral organoids is the absence of microglia and blood vessels, which are thought to emerge during gestational weeks 4-24 (Menassa et al., 2022). Here, we primarily focused on neuroepithelial expansion, which mimics the neurulation stage of development (gestational week 3-4), just before microglia and blood vessels would have developed *in utero.* We conclude that PANX1 is dynamically expressed by multiple cell types in the developing human cerebral cortex. In combination with previous reports from our group and others, this study details the participation of PANX1 in iPSC lineage restriction, co-localizations with key apical membrane proteins and junctional complexes in neuroepithelial rosettes, and PANX1 upregulation and redistribution to TUJ1-expressing neurons within mature human cerebral organoids.

## Ethics Statement

The research performed as a part of this study were approved by the Newfoundland and Labrador Human Research Ethics Board # 2018.201 and # 2014.216.

## Acknowledgements

We thank Dr. Dale Laird for generously providing us with the female iPSCs as well as the anti-human PANX1 C-terminus antibody used in this study. We also thank Dr. Jacqueline Vanderluit for advice regarding brain staining panels and Dr. Matthew Parsons for advice and help with FIJI macros. Finally, we thank Henrietta Odiwa for help with literature searches and blinded organoid scoring.

## Author Contributions

R.J.N. performed the experiments, analyzed the data, and assembled the figures. R.T.F. provided technical assistance. Human fetal brain samples were provided by C.S.M. T.J.B. analyzed the RNAseq data. R.J.N and J.L.E. wrote and edited the manuscript. J.L.E. oversaw the project. All authors reviewed the final version.

## References

Alrefaei AF, Münsterberg AE, Wheeler GN (2020) FZD10 regulates cell proliferation and mediates Wnt1 induced neurogenesis in the developing spinal cord. PLoS One 15:e0219721.

Ang CE et al. (2019) The novel lncRNA. Elife 8.

Baker JC, Beddington RS, Harland RM (1999) Wnt signaling in Xenopus embryos inhibits bmp4 expression and activates neural development. Genes Dev 13:3149–3159.

Beers J, Gulbranson DR, George N, Siniscalchi LI, Jones J, Thomson JA, Chen G (2012) Passaging and colony expansion of human pluripotent stem cells by enzyme-free dissociation in chemically defined culture conditions. Nat Protoc 7:2029–2040.

Bhalla K, Luo Y, Buchan T, Beachem MA, Guzauskas GF, Ladd S, Bratcher SJ, Schroer RJ, Balsamo J, DuPont BR, Lilien J, Srivastava AK (2008) Alterations in CDH15 and KIRREL3 in patients with mild to severe intellectual disability. Am J Hum Genet 83:703–713.

Boassa D, Nguyen P, Hu J, Ellisman MH, Sosinsky GE (2014) Pannexin2 oligomers localize in the membranes of endosomal vesicles in mammalian cells while Pannexin1 channels traffic to the plasma membrane. Front Cell Neurosci 8:468.

Bolte S, Cordelières FP (2006) A guided tour into subcellular colocalization analysis in light microscopy. J Microsc 224:213–232.

Bond SR, Naus CC (2014) The pannexins: past and present. Front Physiol 5:58.

Brandenberger R, Wei H, Zhang S, Lei S, Murage J, Fisk GJ, Li Y, Xu C, Fang R, Guegler K, Rao MS, Mandalam R, Lebkowski J, Stanton LW (2004) Transcriptome characterization elucidates signaling networks that control human ES cell growth and differentiation. Nat Biotechnol 22:707–716.

Carriba P, Davies AM (2017) CD40 is a major regulator of dendrite growth from developing excitatory and inhibitory neurons. Elife 6.

Carriere CH, Wang WX, Sing AD, Fekete A, Jones BE, Yee Y, Ellegood J, Maganti H, Awofala L, Marocha J, Aziz A, Wang LY, Lerch JP, Lefebvre JL (2020) The γ-Protocadherins Regulate the Survival of GABAergic Interneurons during Developmental Cell Death. J Neurosci 40:8652–8668.

Chew L, Añonuevo A, Knock E (2022) Generating Cerebral Organoids from Human Pluripotent Stem Cells. Methods Mol Biol 2389:177–199.

Czermiński JT, Lawrence JB (2020) Silencing Trisomy 21 with XIST in Neural Stem Cells Promotes Neuronal Differentiation. Dev Cell 52:294–308.e293.

Davis LK, Gamazon ER, Kistner-Griffin E, Badner JA, Liu C, Cook EH, Sutcliffe JS, Cox NJ (2012) Loci nominally associated with autism from genome-wide analysis show enrichment of brain expression quantitative trait loci but not lymphoblastoid cell line expression quantitative trait loci. Mol Autism 3:3.

Denaxa M, Kalaitzidou M, Garefalaki A, Achimastou A, Lasrado R, Maes T, Pachnis V (2012) Maturation-promoting activity of SATB1 in MGE-derived cortical interneurons. Cell Rep 2:1351–1362.

Di Lullo E, Kriegstein AR (2017) The use of brain organoids to investigate neural development and disease. Nat Rev Neurosci 18:573–584.

Dobin A, Davis CA, Schlesinger F, Drenkow J, Zaleski C, Jha S, Batut P, Chaisson M, Gingeras TR (2013) STAR: ultrafast universal RNA-seq aligner. Bioinformatics 29:15–21.

Dunn KW, Kamocka MM, McDonald JH (2011) A practical guide to evaluating colocalization in biological microscopy. Am J Physiol Cell Physiol 300:C723–742.

Esseltine JL, Shao Q, Brooks C, Sampson J, Betts DH, Séguin CA, Laird DW (2017) Connexin43 Mutant Patient-Derived Induced Pluripotent Stem Cells Exhibit Altered Differentiation Potential. J Bone Miner Res 32:1368–1385.

Eze UC, Bhaduri A, Haeussler M, Nowakowski TJ, Kriegstein AR (2021) Single-cell atlas of early human brain development highlights heterogeneity of human neuroepithelial cells and early radial glia. Nat Neurosci 24:584–594.

Ferrari ME, Bernis ME, McLeod F, Podpolny M, Coullery RP, Casadei IM, Salinas PC, Rosso SB (2018) Wnt7b signalling through Frizzled-7 receptor promotes dendrite development by coactivating CaMKII and JNK. J Cell Sci 131.

Gyllborg D, Ahmed M, Toledo EM, Theofilopoulos S, Yang S, Ffrench-Constant C, Arenas E (2018) The Matricellular Protein R-Spondin 2 Promotes Midbrain Dopaminergic Neurogenesis and Differentiation. Stem Cell Reports 11:651–664.

Hainz N, Beckmann A, Schubert M, Haase A, Martin U, Tschernig T, Meier C (2018) Human stem cells express pannexins. BMC Res Notes 11:54.

Heo L, Park H, Seok C (2013) GalaxyRefine: Protein structure refinement driven by side-chain repacking. Nucleic Acids Res 41:W384–388.

Hirabayashi Y, Itoh Y, Tabata H, Nakajima K, Akiyama T, Masuyama N, Gotoh Y (2004) The Wnt/beta-catenin pathway directs neuronal differentiation of cortical neural precursor cells. Development 131:2791–2801.

Hou H, Obregon D, Lou D, Ehrhart J, Fernandez F, Silver A, Tan J (2008) Modulation of neuronal differentiation by CD40 isoforms. Biochem Biophys Res Commun 369:641–647.

Islam MM, Zhang CL (2015) TLX: A master regulator for neural stem cell maintenance and neurogenesis. Biochim Biophys Acta 1849:210–216.

Kastriti ME, Stratigi A, Mariatos D, Theodosiou M, Savvaki M, Kavkova M, Theodorakis K, Vidaki M, Zikmund T, Kaiser J, Adameyko I, Karagogeos D (2019) Ablation of CNTN2+ Pyramidal Neurons During Development Results in Defects in Neocortical Size and Axonal Tract Formation. Front Cell Neurosci 13:454.

Kim OK, Nam DE, Hahn YS (2021) The Pannexin 1/Purinergic Receptor P2X4 Pathway Controls the Secretion of MicroRNA-Containing Exosomes by HCV-Infected Hepatocytes. Hepatology 74:3409–3426.

Kriegstein A, Alvarez-Buylla A (2009) The glial nature of embryonic and adult neural stem cells. Annu Rev Neurosci 32:149–184.

Kurosawa H (2007) Methods for inducing embryoid body formation: in vitro differentiation system of embryonic stem cells. J Biosci Bioeng 103:389–398.

Laboratory CSH (2007) Mowial-DABCO stock solution. Cold Spring Harb Protoc.

Lako M, Lindsay S, Bullen P, Wilson DI, Robson SC, Strachan T (1998) A novel mammalian wnt gene, WNT8B, shows brain-restricted expression in early development, with sharply delimited expression boundaries in the developing forebrain. Hum Mol Genet 7:813–822.

Lancaster MA, Knoblich JA (2014) Generation of cerebral organoids from human pluripotent stem cells. Nat Protoc 9:2329–2340.

Lancaster MA, Renner M, Martin CA, Wenzel D, Bicknell LS, Hurles ME, Homfray T, Penninger JM, Jackson AP, Knoblich JA (2013) Cerebral organoids model human brain development and microcephaly. Nature 501:373–379.

Levey DF, Gelernter J, Polimanti R, Zhou H, Cheng Z, Aslan M, Quaden R, Concato J, Radhakrishnan K, Bryois J, Sullivan PF, Stein MB, Program MV (2020) Reproducible Genetic Risk Loci for Anxiety: Results From ∼200,000 Participants in the Million Veteran Program. Am J Psychiatry 177:223–232.

Li S, Qiu F, Xu A, Price SM, Xiang M (2004) Barhl1 regulates migration and survival of cerebellar granule cells by controlling expression of the neurotrophin-3 gene. J Neurosci 24:3104–3114.

Liu S, Rao S, Xu Y, Li J, Huang H, Zhang X, Fu H, Wang Q, Cao H, Baranova A, Jin C, Zhang F (2020) Identifying common genome-wide risk genes for major psychiatric traits. Hum Genet 139:185–198.

Love MI, Huber W, Anders S (2014) Moderated estimation of fold change and dispersion for RNA-seq data with DESeq2. Genome Biol 15:550.

Mancia Leon WR, Spatazza J, Rakela B, Chatterjee A, Pande V, Maniatis T, Hasenstaub AR, Stryker MP, Alvarez-Buylla A (2020) Clustered gamma-protocadherins regulate cortical interneuron programmed cell death. Elife 9.

Masuda T (2017) Contactin-2/TAG-1, active on the front line for three decades. Cell Adh Migr 11:524–531.

Menassa DA, Muntslag TAO, Martin-Estebané M, Barry-Carroll L, Chapman MA, Adorjan I, Tyler T, Turnbull B, Rose-Zerilli MJJ, Nicoll JAR, Krsnik Z, Kostovic I, Gomez-Nicola D (2022) The spatiotemporal dynamics of microglia across the human lifespan. Dev Cell 57:2127–2139.e2126.

Montagne K, Furukawa KS, Taninaka Y, Ngao B, Ushida T (2022) Modulation of the long non-coding RNA Mir155hg by high, but not moderate, hydrostatic pressure in cartilage precursor cells. PLoS One 17:e0275682.

Mordaunt CE, Jianu JM, Laufer BI, Zhu Y, Hwang H, Dunaway KW, Bakulski KM, Feinberg JI, Volk HE, Lyall K, Croen LA, Newschaffer CJ, Ozonoff S, Hertz-Picciotto I, Fallin MD, Schmidt RJ, LaSalle JM (2020) Cord blood DNA methylome in newborns later diagnosed with autism spectrum disorder reflects early dysregulation of neurodevelopmental and X-linked genes. Genome Med 12:88.

Mori T, Buffo A, Götz M (2005) The novel roles of glial cells revisited: the contribution of radial glia and astrocytes to neurogenesis. Curr Top Dev Biol 69:67–99.

Motosugi N, Okada C, Sugiyama A, Kawasaki T, Kimura M, Shiina T, Umezawa A, Akutsu H, Fukuda A (2021) Deletion of lncRNA XACT does not change expression dosage of X-linked genes, but affects differentiation potential in hPSCs. Cell Rep 35:109222.

Neel BL, Nisler CR, Walujkar S, Araya-Secchi R, Sotomayor M (2022) Collective mechanical responses of cadherin-based adhesive junctions as predicted by simulations. Biophys J 121:991–1012.

Noort RJ, Christopher GA, Esseltine JL (2021) Pannexin 1 Influences Lineage Specification of Human iPSCs. Front Cell Dev Biol 9:659397.

Parcerisas A, Pujadas L, Ortega-Gascó A, Perelló-Amorós B, Viais R, Hino K, Figueiro-Silva J, La Torre A, Trullás R, Simó S, Lüders J, Soriano E (2020) NCAM2 Regulates Dendritic and Axonal Differentiation through the Cytoskeletal Proteins MAP2 and 14-3-3. Cereb Cortex 30:3781–3799.

Penuela S, Bhalla R, Nag K, Laird DW (2009) Glycosylation regulates pannexin intermixing and cellular localization. Mol Biol Cell 20:4313–4323.

Penuela S, Lohman AW, Lai W, Gyenis L, Litchfield DW, Isakson BE, Laird DW (2014) Diverse post-translational modifications of the pannexin family of channel-forming proteins. Channels (Austin) 8:124–130.

Penuela S, Bhalla R, Gong XQ, Cowan KN, Celetti SJ, Cowan BJ, Bai D, Shao Q, Laird DW (2007) Pannexin 1 and pannexin 3 are glycoproteins that exhibit many distinct characteristics from the connexin family of gap junction proteins. J Cell Sci 120:3772–3783.

Pluta R, Aragón E, Prescott NA, Ruiz L, Mees RA, Baginski B, Flood JR, Martin-Malpartida P, Massagué J, David Y, Macias MJ (2022) Molecular basis for DNA recognition by the maternal pioneer transcription factor FoxH1. Nat Commun 13:7279.

Prochnow N, Abdulazim A, Kurtenbach S, Wildförster V, Dvoriantchikova G, Hanske J, Petrasch-Parwez E, Shestopalov VI, Dermietzel R, Manahan-Vaughan D, Zoidl G (2012) Pannexin1 stabilizes synaptic plasticity and is needed for learning. PLoS One 7:e51767.

Qu Q, Sun G, Murai K, Ye P, Li W, Asuelime G, Cheung YT, Shi Y (2013) Wnt7a regulates multiple steps of neurogenesis. Mol Cell Biol 33:2551–2559.

Qu Q, Sun G, Li W, Yang S, Ye P, Zhao C, Yu RT, Gage FH, Evans RM, Shi Y (2010) Orphan nuclear receptor TLX activates Wnt/beta-catenin signalling to stimulate neural stem cell proliferation and self-renewal. Nat Cell Biol 12:31–40; sup pp 31-39.

Ray A, Zoidl G, Weickert S, Wahle P, Dermietzel R (2005) Site-specific and developmental expression of pannexin1 in the mouse nervous system. Eur J Neurosci 21:3277–3290.

Renner M, Lancaster MA, Bian S, Choi H, Ku T, Peer A, Chung K, Knoblich JA (2017) Self-organized developmental patterning and differentiation in cerebral organoids. EMBO J 36:1316–1329.

Rosso SB, Sussman D, Wynshaw-Boris A, Salinas PC (2005) Wnt signaling through Dishevelled, Rac and JNK regulates dendritic development. Nat Neurosci 8:34–42.

Salik D, Hadj-Rabia S, Hohl D, Vahidnezhad H, Youssefian L, Rakosi A, Dangoisse C, Marangoni M, Vilain C, Smits G (2022) Evaluation of neurodevelopmental symptoms in 10 cases of neonatal ichthyosis and sclerosing cholangitis syndrome. Pediatr Dermatol 39:590–593.

Sanchez-Arias JC, Candlish RC, van der Slagt E, Swayne LA (2020) Pannexin 1 Regulates Dendritic Protrusion Dynamics in Immature Cortical Neurons. eNeuro 7.

Sang Q et al. (2019) A pannexin 1 channelopathy causes human oocyte death. Sci Transl Med 11.

Sanmaneechai O et al. (2015) Genotype-phenotype characteristics and baseline natural history of heritable neuropathies caused by mutations in the MPZ gene. Brain 138:3180–3192.

Sayedyahossein S, Huang K, Li Z, Zhang C, Kozlov AM, Johnston D, Nouri-Nejad D, Dagnino L, Betts DH, Sacks DB, Penuela S (2021) Pannexin 1 binds β-catenin to modulate melanoma cell growth and metabolism. J Biol Chem:100478.

Schindelin J, Arganda-Carreras I, Frise E, Kaynig V, Longair M, Pietzsch T, Preibisch S, Rueden C, Saalfeld S, Schmid B, Tinevez JY, White DJ, Hartenstein V, Eliceiri K, Tomancak P, Cardona A (2012) Fiji: an open-source platform for biological-image analysis. Nat Methods 9:676–682.

Seo JH, Dalal MS, Contreras JE (2021) Pannexin-1 Channels as Mediators of Neuroinflammation. Int J Mol Sci 22.

Shao Q, Lindstrom K, Shi R, Kelly J, Schroeder A, Juusola J, Levine KL, Esseltine JL, Penuela S, Jackson MF, Laird DW (2016) A Germline Variant in the PANX1 Gene Has Reduced Channel Function and Is Associated with Multisystem Dysfunction. J Biol Chem 291:12432–12443.

Sharma K, Choi SY, Zhang Y, Nieland TJ, Long S, Li M, Huganir RL (2013) High-throughput genetic screen for synaptogenic factors: identification of LRP6 as critical for excitatory synapse development. Cell Rep 5:1330–1341.

Sheng L, Leshchyns’ka I, Sytnyk V (2015) Neural cell adhesion molecule 2 promotes the formation of filopodia and neurite branching by inducing submembrane increases in Ca2+ levels. J Neurosci 35:1739–1752.

Shum MG, Shao Q, Lajoie P, Laird DW (2019) Destination and consequences of Panx1 and mutant expression in polarized MDCK cells. Exp Cell Res 381:235–247.

Speevak MD, Farrell SA (2013) Charcot-Marie-Tooth 1B caused by expansion of a familial myelin protein zero (MPZ) gene duplication. Eur J Med Genet 56:566–569.

Takeuchi C, Ishikawa M, Sawano T, Shin Y, Mizuta N, Hasegawa S, Tanaka R, Tsuboi Y, Nakatani J, Sugiura H, Yamagata K, Tanaka H (2020) Dendritic Spine Density is Increased in Arcadlin-deleted Mouse Hippocampus. Neuroscience 442:296–310.

Tran TM, Sherwood JK, Doolittle MJ, Sathler MF, Hofmann F, Stone-Roy LM, Kim S (2021) Loss of cGMP-dependent protein kinase II alters ultrasonic vocalizations in mice, a model for speech impairment in human microdeletion 4q21 syndrome. Neurosci Lett 759:136048.

Turmel P, Dufresne J, Hermo L, Smith CE, Penuela S, Laird DW, Cyr DG (2011) Characterization of pannexin1 and pannexin3 and their regulation by androgens in the male reproductive tract of the adult rat. Mol Reprod Dev 78:124–138.

Wang C, Pan YH, Wang Y, Blatt G, Yuan XB (2019) Segregated expressions of autism risk genes Cdh11 and Cdh9 in autism-relevant regions of developing cerebellum. Mol Brain 12:40.

Wang DY, Luo AF, Bai QR, Gong XL, Zheng Y, Shen Q, Hu XL, Wang XM (2020a) VCAM1 Labels a Subpopulation of Neural Stem Cells in the Adult Hippocampus and Contributes to Spatial Memory. Stem Cell Reports 14:1093–1106.

Wang Y, Kerrisk Campbell M, Tom I, Foreman O, Hanson JE, Sheng M (2020b) PCDH7 interacts with GluN1 and regulates dendritic spine morphology and synaptic function. Sci Rep 10:10951.

Watanabe K, Ueno M, Kamiya D, Nishiyama A, Matsumura M, Wataya T, Takahashi JB, Nishikawa S, Muguruma K, Sasai Y (2007) A ROCK inhibitor permits survival of dissociated human embryonic stem cells. Nat Biotechnol 25:681–686.

Wicki-Stordeur LE, Swayne LA (2013) Panx1 regulates neural stem and progenitor cell behaviours associated with cytoskeletal dynamics and interacts with multiple cytoskeletal elements. Cell Commun Signal 11:62.

Wicki-Stordeur LE, Dzugalo AD, Swansburg RM, Suits JM, Swayne LA (2012) Pannexin 1 regulates postnatal neural stem and progenitor cell proliferation. Neural Dev 7:11.

Wicki-Stordeur LE, Sanchez-Arias JC, Dhaliwal J, Carmona-Wagner EO, Shestopalov VI, Lagace DC, Swayne LA (2016) Pannexin 1 Differentially Affects Neural Precursor Cell Maintenance in the Ventricular Zone and Peri-Infarct Cortex. J Neurosci 36:1203–1210.

Wilson PG, Stice SS (2006) Development and differentiation of neural rosettes derived from human embryonic stem cells. Stem Cell Rev 2:67–77.

Wingett SW, Andrews S (2018) FastQ Screen: A tool for multi-genome mapping and quality control. F1000Res 7:1338.

Xiao Y, Hu M, Lin Q, Zhang T, Li S, Shu L, Song X, Xu X, Meng W, Li X, Xu H, Mo X (2023) Dopey2 and Pcdh7 orchestrate the development of embryonic neural stem cells/ progenitors in zebrafish. iScience 26:106273.

Yan C, Meng Y, Yang J, Chen J, Jiang W (2023) Translational landscape in human early neural fate determination. Development 150.

Yan Y, Shin S, Jha BS, Liu Q, Sheng J, Li F, Zhan M, Davis J, Bharti K, Zeng X, Rao M, Malik N, Vemuri MC (2013) Efficient and rapid derivation of primitive neural stem cells and generation of brain subtype neurons from human pluripotent stem cells. Stem Cells Transl Med 2:862–870.

Yang SL, Yang M, Herrlinger S, Liang C, Lai F, Chen JF (2015) MiR-302/367 regulate neural progenitor proliferation, differentiation timing, and survival in neurulation. Dev Biol 408:140–150.

Yasuda S, Tanaka H, Sugiura H, Okamura K, Sakaguchi T, Tran U, Takemiya T, Mizoguchi A, Yagita Y, Sakurai T, De Robertis EM, Yamagata K (2007) Activity-induced protocadherin arcadlin regulates dendritic spine number by triggering N-cadherin endocytosis via TAO2beta and p38 MAP kinases. Neuron 56:456–471.

Young MD, Wakefield MJ, Smyth GK, Oshlack A (2010) Gene ontology analysis for RNA-seq: accounting for selection bias. Genome Biol 11:R14.

Zandi PP, Belmonte PL, Willour VL, Goes FS, Badner JA, Simpson SG, Gershon ES, McMahon FJ, DePaulo JR, Potash JB, Group BDP, Consortium NIoMHGIBD (2008) Association study of Wnt signaling pathway genes in bipolar disorder. Arch Gen Psychiatry 65:785–793.

Zappalà A, Li Volti G, Serapide MF, Pellitteri R, Falchi M, La Delia F, Cicirata V, Cicirata F (2007) Expression of pannexin2 protein in healthy and ischemized brain of adult rats. Neuroscience 148:653–667.

Zarrei M et al. (2019) A large data resource of genomic copy number variation across neurodevelopmental disorders. NPJ Genom Med 4:26.

Zhao Z, Zhang D, Yang F, Xu M, Zhao S, Pan T, Liu C, Liu Y, Wu Q, Tu Q, Zhou P, Li R, Kang J, Zhu L, Gao F, Wang Y, Xu Z (2022) Evolutionarily conservative and non-conservative regulatory networks during primate interneuron development revealed by single-cell RNA and ATAC sequencing. Cell Res 32:425–436.

Zhu L, Gomez-Duran A, Saretzki G, Jin S, Tilgner K, Melguizo-Sanchis D, Anyfantis G, Al-Aama J, Vallier L, Chinnery P, Lako M, Armstrong L (2016) The mitochondrial protein CHCHD2 primes the differentiation potential of human induced pluripotent stem cells to neuroectodermal lineages. J Cell Biol 215:187–202.

